# Integrative genomics reveals paths to sex dimorphism in *Salix purpurea* L.

**DOI:** 10.1101/2021.03.09.434562

**Authors:** Brennan Hyden, Craig H. Carlson, Fred E. Gouker, Jeremy Schmutz, Kerrie Barry, Anna Lipzen, Aditi Sharma, Laura Sandor, Gerald A. Tuskan, Guanqiao Feng, Matthew S. Olson, Stephen P. DiFazio, Lawrence B. Smart

## Abstract

Sex dimorphism and gene expression were studied in developing catkins in 159 F_2_ individuals from the bioenergy crop *Salix purpurea*, and potential mechanisms and pathways for regulating sex development were explored. Differential expression, eQTL, bisulfite sequencing, and network analysis were used to characterize sex dimorphism, detect candidate master regulator genes, and identify pathways through which the sex determination region (SDR) may mediate sex dimorphism. Eleven genes are presented as candidates for master regulators of sex, supported by gene expression and network analyses. These include genes putatively involved in hormone signaling, epigenetic modification, and regulation of transcription. eQTL analysis revealed a suite of transcription factors and genes involved in secondary metabolism and floral development that were predicted to be under direct control of the sex determination region. Furthermore, data from bisulfite sequencing and small RNA sequencing revealed strong differences in expression between males and females that would implicate both of these processes in sex dimorphism pathways. These data indicate that the mechanism of sex determination in *Salix purpurea* is likely different from that observed in the related genus *Populus*. This further demonstrates the dynamic nature of SDRs in plants, which involves a multitude of mechanisms of sex determination and a high rate of turnover.

## Introduction

Major progress has been made in recent years in identifying the master regulators of sex determination in plants, but less is known about the transcriptional networks that account for sex dimorphism. Transcription factors^1^ (*Asparagus*), small RNAs^2^ (*Diospyros*) and cytokinin response regulators (*Actinida* and *Populus*)^3,4^ have all been explicated as mechanisms of sex determination in angiosperms. Development of separate sexes, however, requires that the genes controlling sex determination regulate the transcription of many genes and metabolic pathways in order to coordinate development of sex-specific characteristics, such as gametes and floral morphology. With the notable exception of *Diospyros*^*5*^, little is known about the metabolic pathways that are regulated by these sex-determination genes and the resulting transcriptome-wide expression differences.

Evidence from multiple angiosperm taxa, as well as leading sex determination models, suggest that in plants, sex is controlled by one or two regulatory factors, termed “sex determination” or “master regulator” genes^1,2,4,6,7^. These factors in turn lead to primary sex dimorphisms (androecium and gynoecium development) as well as secondary sex dimorphisms, such as floral volatile profiles, pigmentation, floral phenology, and organ morphology, and often involve both sex linked and autosomal genes^8^. While primary sex dimorphisms are under direct control of the sex determination gene(s), secondary sex dimorphisms may be either under the control of sex - linked genes, or under direct control of the sex-determining genes themselves^7,8^. Because their expression in the opposite sex may be deleterious, linkage between sex determination genes and genes controlling secondary sexual dimorphisms may be favored by natural selection. Such genes are termed “sexually antagonistic”^9^. As a result, it can be challenging to discriminate among sex determination genes and other sex-linked genes that influence sex dimorphisms.

The maintenance of separate sexes requires that the factors controlling sex determination, along with linked sexually antagonistic genes controlling sex dimorphisms, reside on heterogametic sex chromosomes, where typically one sex is heterogametic and the other homogametic^6^. Two heterogametic systems, XY and ZW, have been observed in angiosperms. The XY system, where the male is the heterogametic sex, tends to be more prevalent, and is found in 84.7% of dioecious angiosperm species, including *Asparagus officinalis, Carica papaya, Diospyros*, and *Phoenix dactylifera*. In the less common ZW system, females are heterogametic, as in *Fragaria* and *Silene otitis*^*10,11*^. The Salicaceae family is of particular interest for use as a model in understanding the evolution of sex chromosomes and sex determination in plants, because the family is almost exclusively dioecious, yet the sex determination region (SDR) has been remarkably dynamic^4,12,13^. The Salicaceae family exhibits both ZW and XY heterogametic systems and SDRs that are localized in different chromosomes across species. The SDR in *S. purpurea* has been localized to a 6.73 Mb pericentromeric region on chr15W, which includes approximately 2 Mb of sequence that is not present in the corresponding region of chr15Z^14^. Other *Salix* spp. in the section *Vetrix* also have a chr15 ZW SDR, including *S. viminalis* and *S. suchowensis*^*15,16*^, whereas *S. nigra* in the section *Salix* has a chr07 XY system^12^. In contrast, most *Populus* species have chr19 SDRs, including XY in *P. trichocarpa*^*17*^ and ZW in *P. alba*^*18*^ but, exceptionally, *P. euphratica* exhibits a chr14 XY SDR^19^. This indicates that sex determination has a complex evolutionary history in the Salicaceae, and that the SDR has shifted chromosomes as well as heterogametic systems, either through translocation, the rise of an entirely new locus becoming sexually antagonistic, or that there have been several independent origins of dioecy.

The Salicaceae family contains many species of economic importance in the genera *Populus* and *Salix*. Shrub willows (*Salix* section *Vetrix*) in particular, are grown throughout North America and Eurasia for bioenergy and bio-products^20^. Despite its commonality across the Salicaceae family, dioecy presents a challenge for breeding efforts and the cultivation of shrub willow, with sex showing linkage to biomass related traits, such as leaf area^21^ and catkins showing distinct phenology and secondary metabolite profiles between sexes, affecting pollinator and pest attraction^22,23^. There is a strong interest in understanding the genetic mechanisms controlling sex determination in *Salix*, along with the gene pathways involved in sex dimorphism, in order to advance current breeding efforts and genetic studies to improve *Salix* as a bioenergy crop. Nevertheless, there is still relatively little data characterizing sex-determination genes and sex dimorphism pathways in *Salix*, despite substantial research and identification of putative master regulators of sex in the related genus *Populus*^*4,18,24,25*^. Moreover, willows are typically insect pollinated, whereas members of the genus *Populus* are wind pollinated and show little evidence of sex dimorphism in vegetative traits^26^ which may point to unique pathways and genes involved in sex dimorphism and underscores the need for sex determination research that is specific to *Salix*.

A previous study of transcriptomic data in *Salix* identified differentially expressed genes associated with sex in shoot tips containing floral primordia^27^. It is hypothesized that sexually dimorphic genes are regulated through complex pathways that are ultimately controlled by one or more elements in the SDR, most likely via transcription factors, plant hormones, and/or small RNAs. This would be consistent with characterized sex determination systems in plants^1–4^. Elements in the SDR controlling sex determination are termed “master regulator genes” and likely control several top-level regulatory genes that may or may not be located in the SDR, which in turn regulate both primary and secondary sex characteristics through intermediate metabolic pathways, as described by Feng et al., 2020^11^. Unfortunately, identification of master regulator genes in *Salix purpurea* is complicated by an SDR that comprises nearly 40% of chr15W and contains 488 linked genes, including repetitive regions, and tandem duplications^14^, requiring the use of transcriptomic data and co-expression analyses to characterize candidate genes.

This study captures the transcriptome-wide primary and secondary sex dimorphism profile in emerging inflorescences, which contain hundreds of achlamydeous flowers across a range of developmental time points, in addition to exhibiting distinct terpenoid and phenylpropanoid profiles leading to pigmentation and volatile organic compound emission dimorpism^22^. Using eQTL analysis, we associated the expression levels of differentially expressed genes in catkins with genomic loci. We identified multiple genes associated in *trans* with the SDR that could serve as top-level regulators of primary and secondary floral sex dimorphisms under direct control by master regulator genes, as well as enriched pathways predicted to serve as intermediate pathways involved in sex dimorphism. Furthermore, based on these multi-omics results, six gene families are presented as candidates for the master regulators of sex: homologs of Arabidopsis *GATA15, ARR17, AGO4, DRB1*, three genes coding hypothetical proteins, and a CCHC zinc finger. Characterizing the master regulator genes and the mechanisms of sex determination in willow provides insight into the complex evolution of dioecy in the Salicaceae. These data represent one of the most comprehensive studies of sex dimorphism expression in a dioecious crop plant, incorporating RNA expression, genotyping-by-sequencing (GBS), differential methylation, and small RNA expression, and provide a valuable addition to the nascent body of knowledge on sex determination mechanisms in crop plants.

## Results

### Differential Expression Analysis

Principal component analysis showed a clear separation of male and female genotypes based on transcriptome-wide expression (Fig. S1). A total of 36,518 gene models, including alternative transcripts, were expressed in floral tissue across the 159 samples, with 24,074 (66%) showing significant differential expression between males (M) and females (F) (FDR ≤ 0.05) (Fig. 1). There were 4,676 genes with log_2_ M:F ≥ 1 (male upregulated), while only 3,247 genes with log_2_ M:F ≤ −1 (female upregulated).

**Fig 1.**
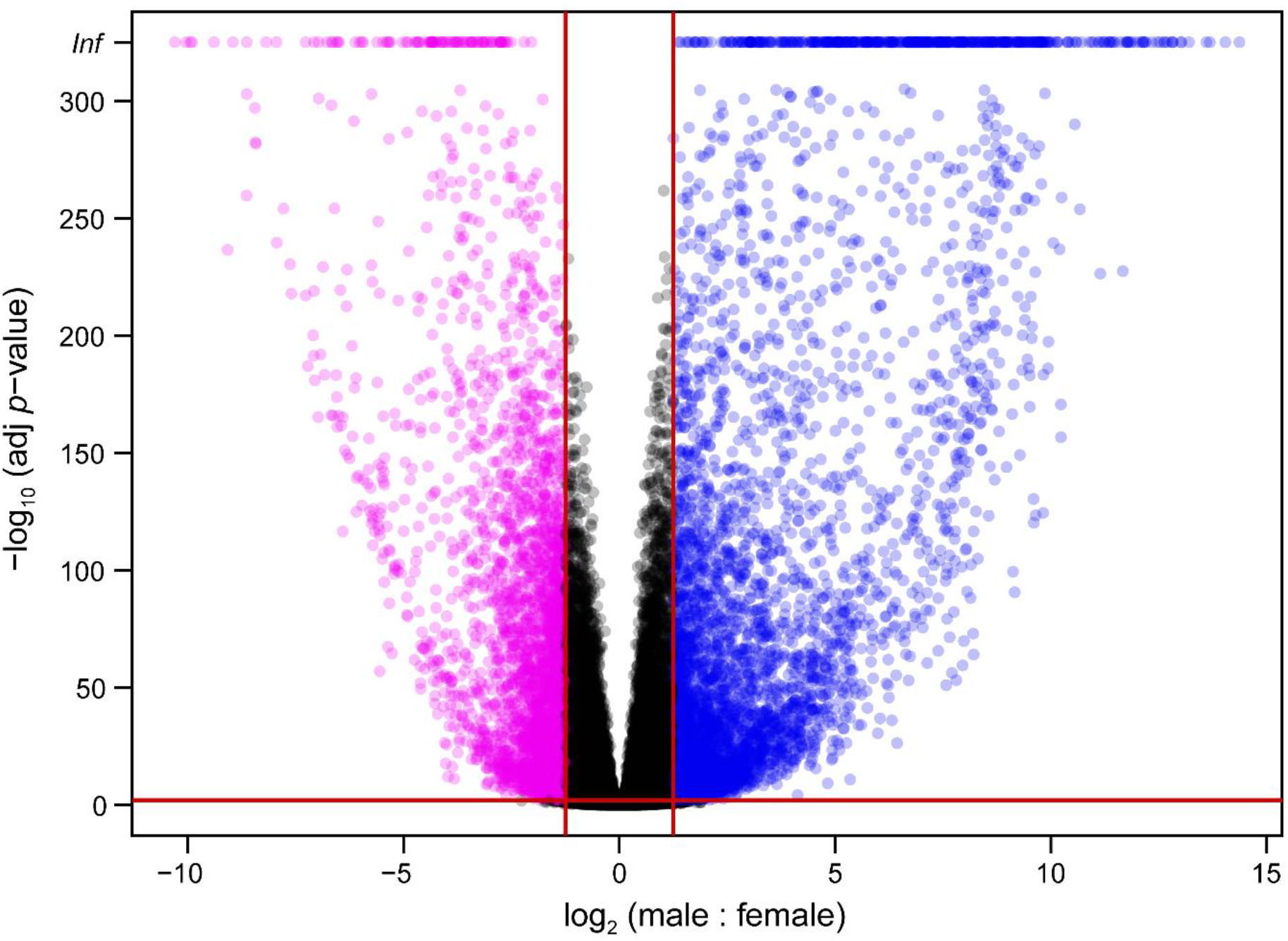
Volcano plot showing the log_2_ differential gene expression between males and females compared to FDR adjusted p-value. Blue points indicate significantly male upregulated genes while magenta points indicate those upregulated in females.

Gene Ontology enrichment was performed on the male and female differentially expressed (DE) genes showing at least four-fold expression differences between sexes (Tables S1-S4). Gene Ontology enrichment for the male upregulated genes showed a significant overrepresentation of 76 terms, including pollen and anther development, male gamete development, the terpenoid biosynthesis pathway, and cytokinin metabolism (Table S1). The male upregulated gene set showed an underrepresentation for 71 terms, including terms relating to transcription and RNA regulation, splicing, and modification (Table S2). Among the female upregulated genes, there was an overrepresentation of RNA transcription and metabolism and shoot and organ development (Table S3), and an underrepresentation of 47 terms including cell metabolism and biosynthesis (Table S4).

### Small RNA identification and analysis

A total of 266,272 small RNA (smRNA) loci were identified, 146,807 of which contained smRNAs in the range of 20-25 nucleotides, the canonical length of siRNAs and miRNAs. Forty-five miRNA precursor loci representing 34 unique final miRNAs were predicted based on putative stem-loop structures consistent with miRNA biosynthesis (Table 1, Fig. 2, Table S5, Fig. S2). Forty of the 45 loci had identity with known and annotated plant smRNAs in the PmiREN database (www.pmiren.com). psRNATarget (plantgrn.noble.org/psRNATarget) was used to identify putative target genes for all miRNAs. Differential expression analysis on the miRNA loci revealed that 25 were significantly differentially expressed (FDR ≤ 0.05), including 18 with greater expression in males and seven with greater expression in females. Among the miRNA loci were two male upregulated copies of miR172, which targets the MADS-box gene *APETALA2* in Arabidopsis^*28*^ and four male upregulated copies of miR156, one of which is located in the SDR.

**Table 1.**
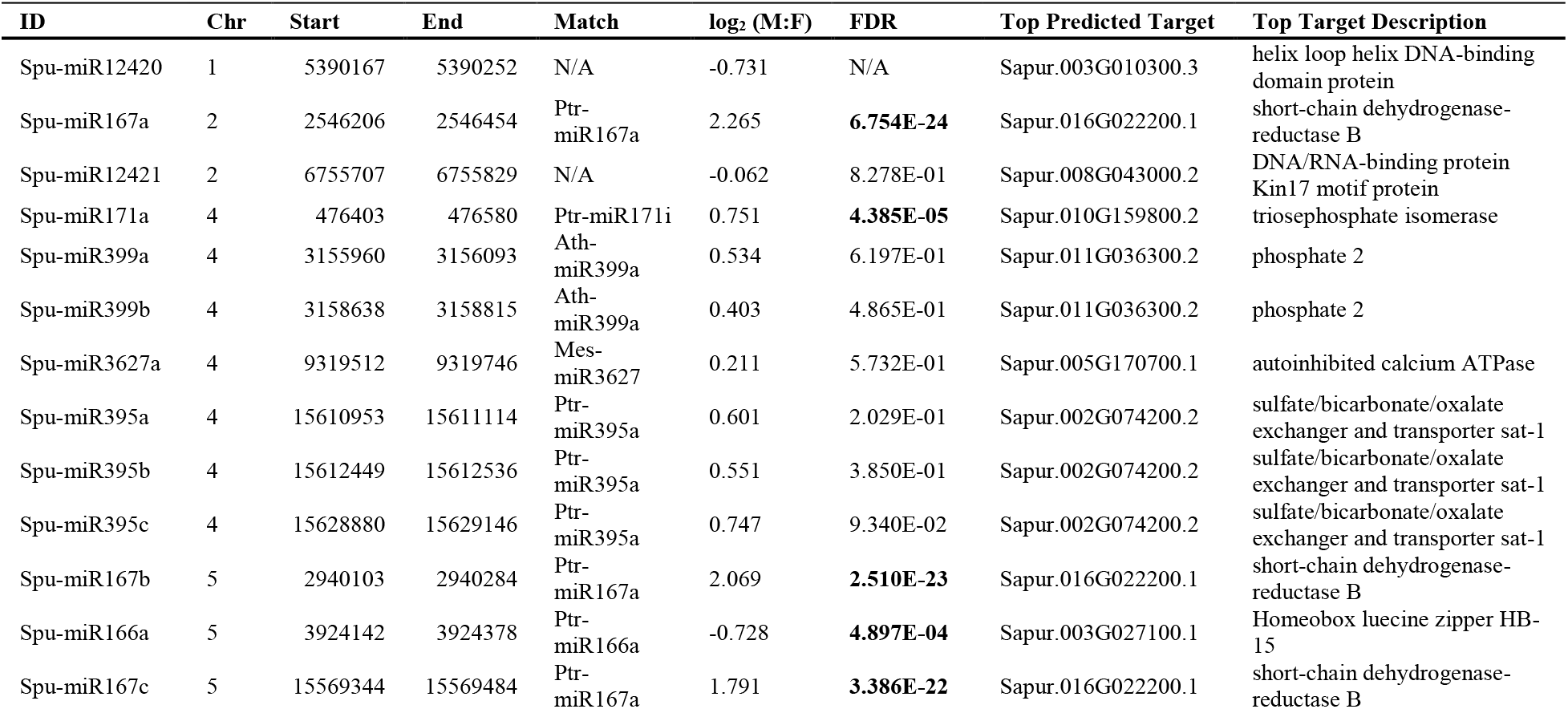
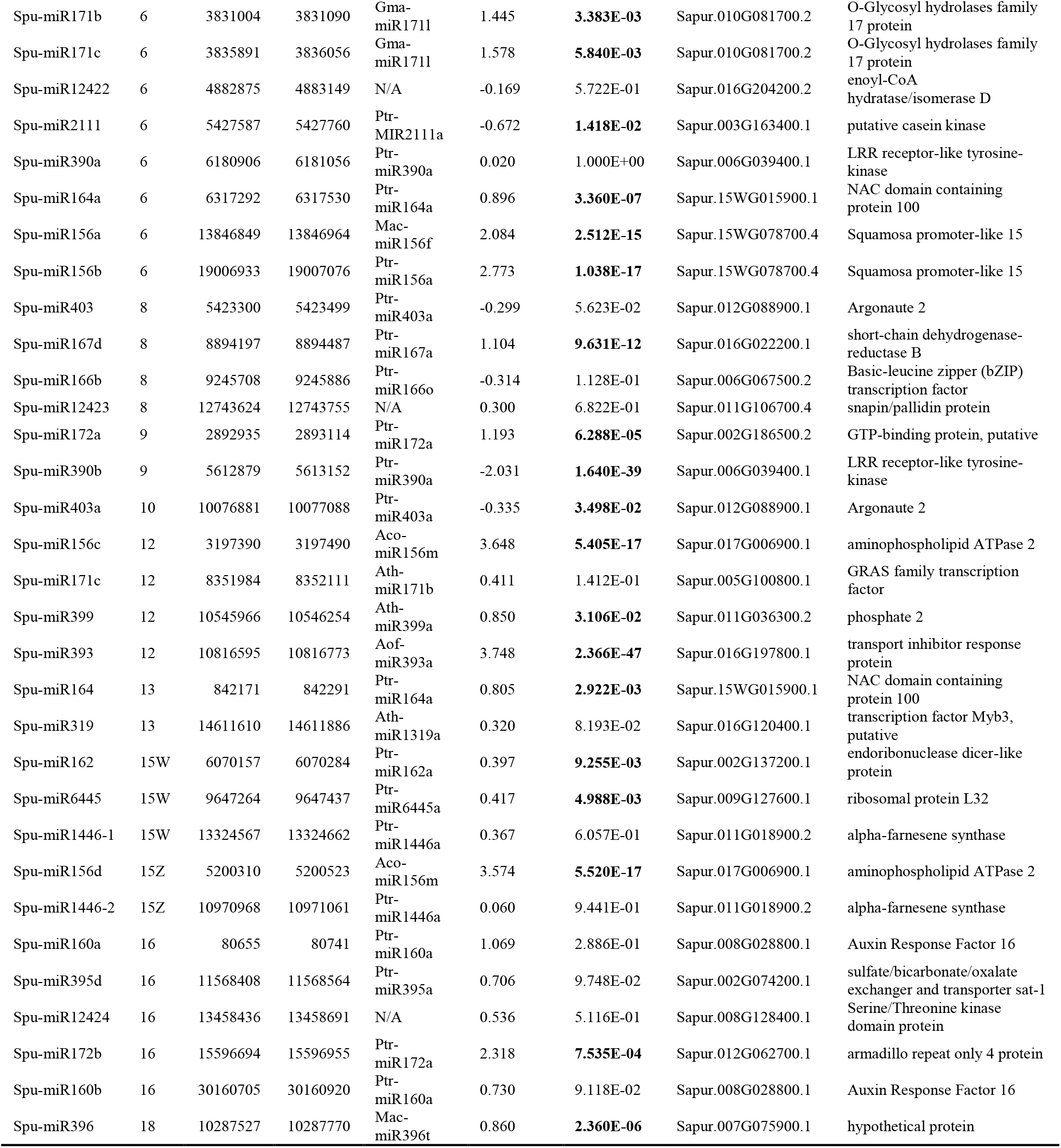
Forty-five miRNA loci identified and their respective PmiREN database matches with known plant small RNAs as well as the top predicted hit in the *S. purpurea* genome using psRNATarget. Negative log_2_-fold change values are female upregulated and positive values are male upregulated. Those with significant FDR values are in bold.

**Fig 2.**
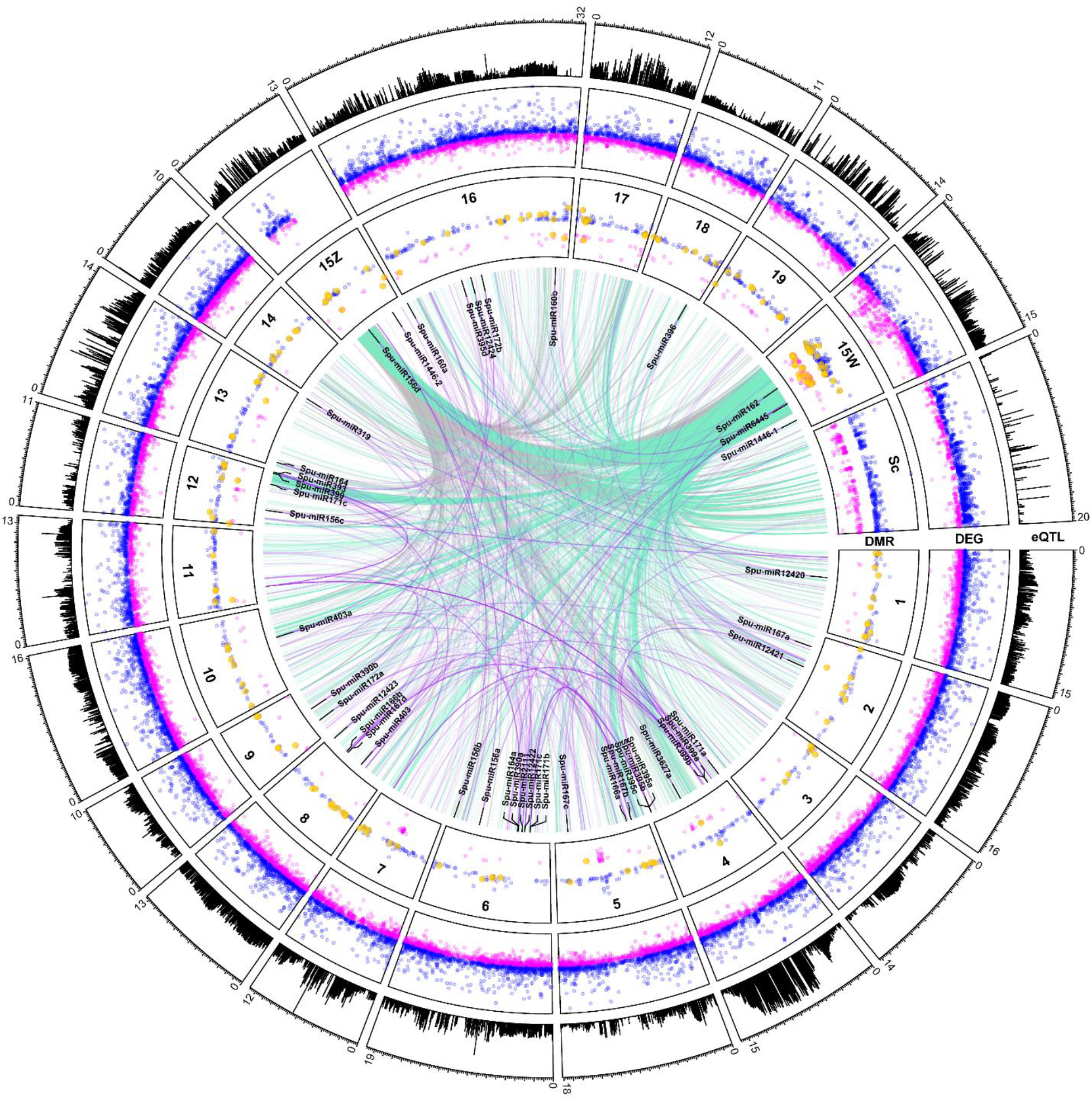
Circos plot showing total eQTLs per locus on the outer track (range 0 to 550), mapping sites of all differentially methylated regions in the middle track (male upregulated in blue, female upregulated in magenta, DMRs in putative gene promoters in gold), and mapping sites of all differentially expressed genes in the inner track. Associations between gene expression and polymorphisms in the Z-SDR are shown in grey while associations with the W-SDR are shown in aquamarine. The top ten predicted target sites for each miRNA are shown in purple.

### Bisulfite Sequencing and Analysis

A total of 604,150 methylated regions were identified using DMRfinder after combining nearby methylated sites. Of these sites, 2,018 show differential methylation, including 1,465 with greater methylation in males and 553 with greater methylation in females (Table S6). 170 genes were identified with differentially methylated sites in their putative promoter regions (between 1 and 1000 bp ahead of transcription start site) (Table S7).

### WGCNA Network Analysis and GO Enrichment

WGCNA network analysis was performed to explore pathways that may be involved in floral sex dimorphism and development. Twenty-four modules were generated based on gene expression similarities (Fig. 3, Table 2). 17,953 genes were assigned to modules, accounting for 49.8% of all expressed genes and alternative transcripts. The remaining unassigned genes were placed in the “grey” module. Three modules accounted for the majority of assigned genes: the “purple”, “cyan” and “brown” modules. The “purple” module was strongly correlated to the female sex (r^2^=0.97) and captured 30.2% of all female expressed genes (Table 2), this module likely represents the genes involved in primary and secondary female sex dimorphism. In the “purple” module 17 GO terms were enriched, including RNA and nucleic acid metabolism and regulation, photosynthesis, and phenylpropanoid metabolism (Table S8). This is consistent with the female differential expression of the majority of genes in this module (Table S3) and supports the contention that this module may be responsible for female sex-dimorphic traits. The cyan module was the most male correlated (r^2^=0.92) and included 45.5% of all male upregulated genes likely representing the genes in primary and secondary male sex dimorphism pathways. The “cyan” module contained 230 significantly enriched GO terms, notably multiple terms related to pollen development (Table S9). The “brown” module consisted mostly of genes not showing differential expression, and probably represents gene pathways involved in basic cell and biological processes.

**Table 2:**
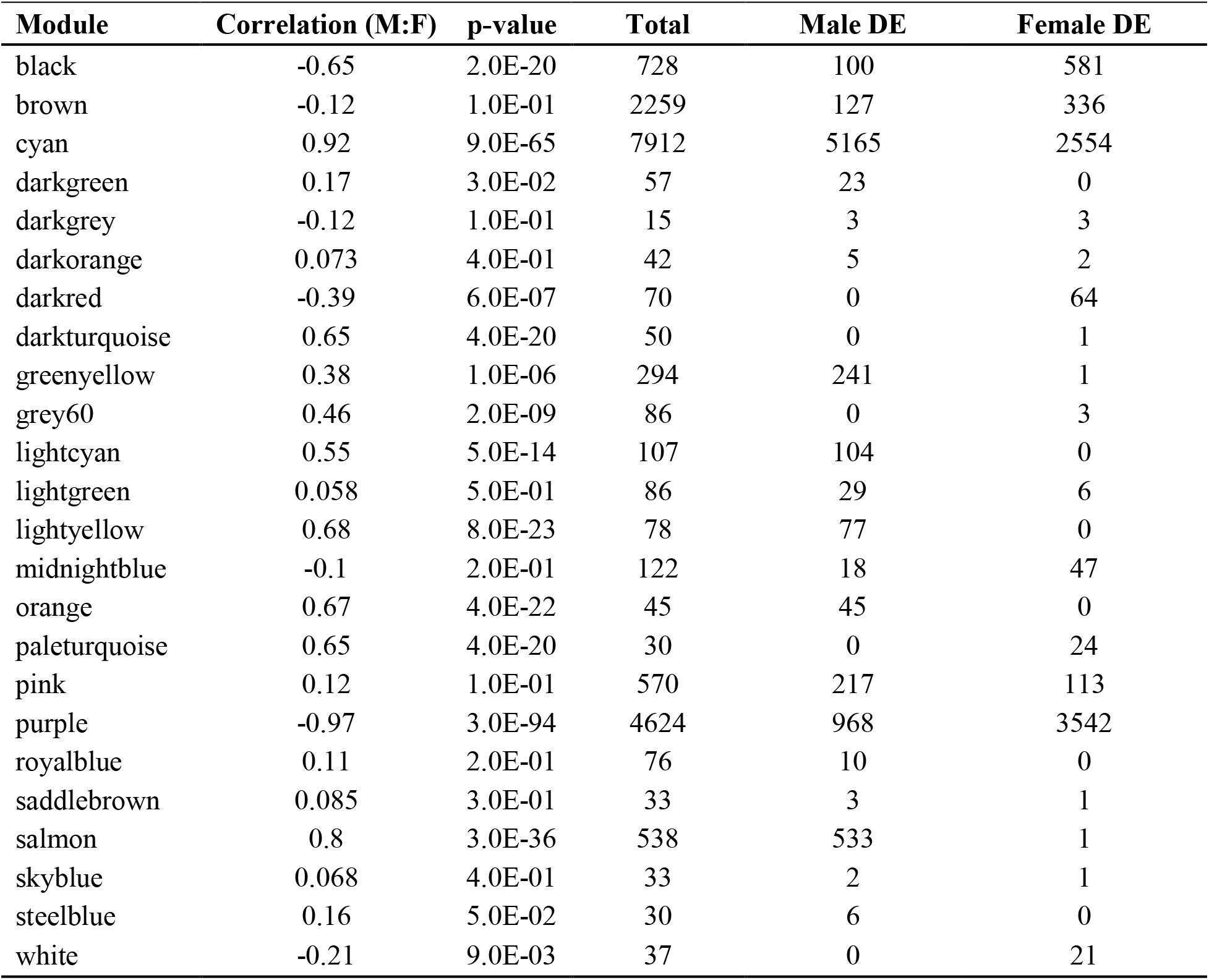
All WGCNA modules with the total number of genes belonging to each, along with the number of differentially expressed genes as determined through DESEQ2 analysis; the grey module contains all genes that were unable to be assigned to a specific module.

**Figure 3.**
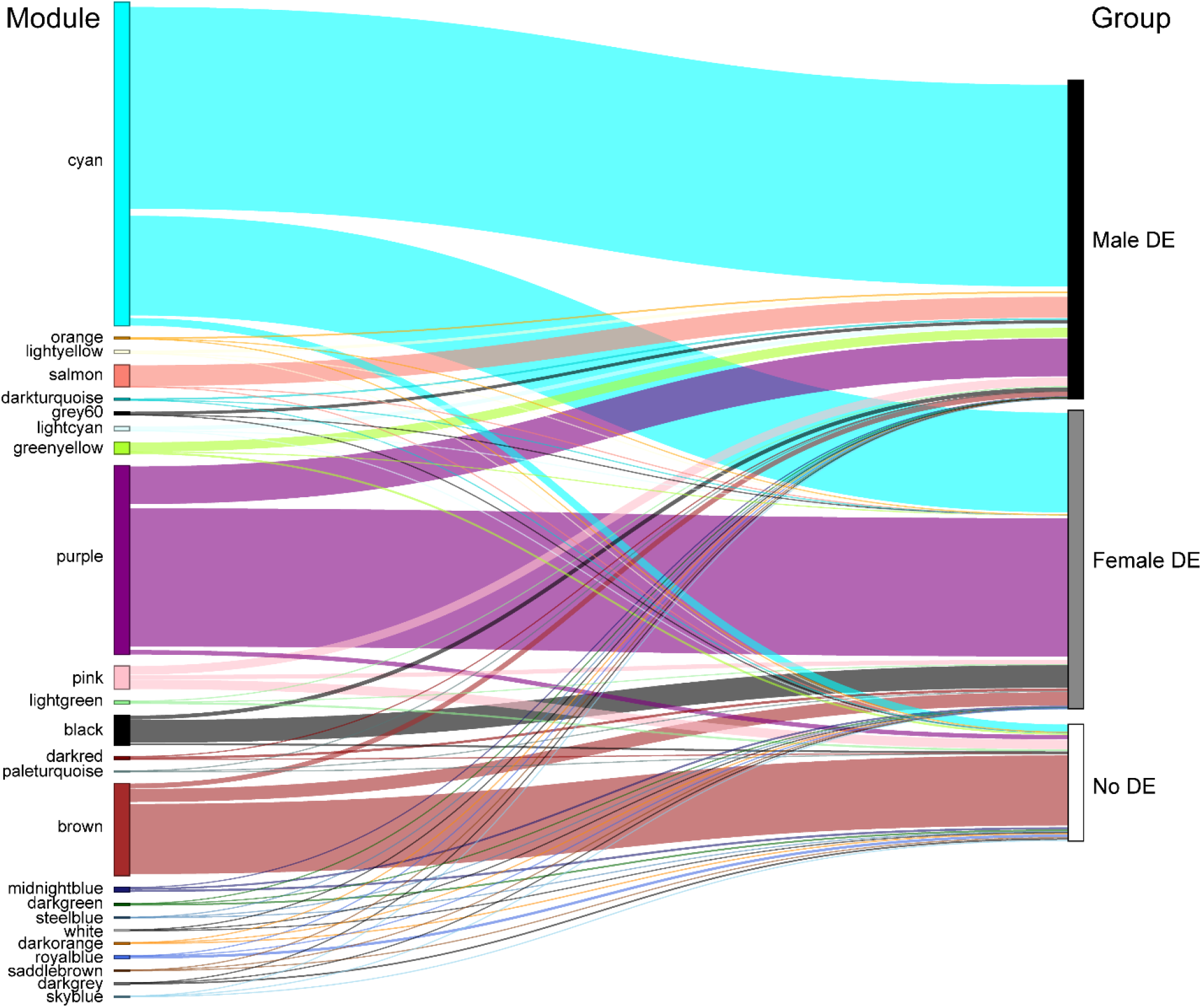
Sankey plot showing the relative size of each WGCNA module and the proportion of male, female, and non-upregulated genes. The unassigned genes that were placed in the “grey” module are not shown.

### eQTL Analysis

A total of 1,381,813 *cis* eQTL and 811,499 *trans* eQTL (FDR ≤ 0.05) were identified after accounting for covariates. Notably, there appears to be an eQTL “hotspot” on chr07, to which the expression levels of 550 genes are associated (Fig. 2) The exceptional number of genomewide eQTL at this locus suggests that it may have a major role in regulating sex dimorphic gene expression in catkin tissue. Expression levels of 2,127 genes were found to be associated with polymorphisms in the SDR, of which 1,686 were *trans* (Fig. 2). To identify top level regulatory and intermediate pathway genes, 2,127 genes with eQTL in the SDR were subset for genes with involvement in either secondary metabolism (22), hormone signaling (16), RNA splicing and regulation (4) or transcription (96) (Table S10, Fig. S3). Of the 96 genome-wide transcription factors found to have SDR eQTL, 70 were differentially expressed in either males or females. Because these transcription factors include genes relating to floral development, phenylpropanoid production, and cytokinin signaling, they are candidates for top-level regulatory genes that may regulate further downstream expression. Fourteen MADS-box and floral development genes, 15 phenylpropanoid pathway genes, and five terpenoid pathway genes were also found to have eQTL in the SDR (Table S10), representing candidates for intermediate pathway genes directly responsible for dimorphisms in floral morphology, pigmentation, and volatile and secondary metabolite profiles.

### Candidate Master Regulator Gene Identification

We identified eleven gene that are strong candidates as master regulators of sex using the following criteria: (1) presence on chr15W and absence from chr15Z; (2) a significant log _2_ M:F < −1; (3) presence in the female correlated “purple” WGCNA module, and (4) gene annotation either consistent with a possible floral sex dimorphism pathway or of unknown function. Genes meeting all four of these criteria are expected to be present only in females and have expression levels and module membership that would implicate them in sex dimorphism. Four copies of *ARR17*, a truncated *AGO4* gene, *DRB1, GATA15*, a CCHC zinc finger nuclease, and three genes coding hypothetical proteins met these criteria and were identified as candidate master regulator genes (Table 3).

**Table 3.** Eleven candidate master regulator genes and their log_2_ male:female (M:F) differential expression values and *Arabidopsis thaliana* homologies.

| Gene ID | $\log_2$ (M:F) | FDR | Arabidopsis homolog | Arabidopsis gene name | Description |
| --- | --- | --- | --- | --- | --- |
| Sapur.15WG062800.1 | -7.491 | 2.23E-250 | AT3G06740.1 | *GATA15* | GATA transcription factor 15 |
| Sapur.15WG068800.1 | -6.282 | 5.82E-167 | AT2G15180.1 | - | Zinc knuckle (CCHC-type) family protein |
| Sapur.15WG073500.1 | -3.973 | 1.67E-60 | AT3G56380.1 | *ARR17* | response regulator 17 |
| Sapur.15WG073900.1 | -2.802 | 1.93E-25 | AT3G56380.1 | *ARR17* | response regulator 17 |
| Sapur.15WG074000.1 | -4.026 | 3.82E-62 | AT3G56380.1 | *ARR17* | response regulator 17 |
| Sapur.15WG074300.1 | -1.845 | 5.68E-13 | AT1G09700.1 | *DRB1* | dsRNA-binding domain-like protein |
| Sapur.15WG074400.1 | -7.842 | 3.99E-263 | AT2G27040.1 | *AGO4* | Argonaute family protein |
| Sapur.15WG074900.1 | -1.305 | 1.06E-25 | - | - | - |
| Sapur.15WG075200.1 | -4.004 | 3.94E-61 | AT3G56380.1 | *ARR17* | response regulator 17 |
| Sapur.15WG075300.2 | -3.534 | 4.02E-52 | AT1G77270.1 | - | hypothetical protein |
| Sapur.15WG075700.1 | -3.840 | 2.71E-53 | - | - | - |

## Discussion

We have used a combination of differential expression, co-expression, and eQTL analyses to identify genes that are candidate master and top-level regulators of sex expression in *S. purpurea* floral tissue.

### RNA-Seq and Differential Expression

Total RNA-Seq and small RNA-Seq captured the unique sex-specific transcriptomic profiles during catkin development, after floral meristem differentiation and prior to maturation of any stamens or pistils. Within a single maturing catkin, there are hundreds of individual flowers across a range of developmental stages, resulting in pooled expression data from across floral development time points as well as tissue types (i.e., anthers/pistils, floral bracts, and peduncles). In addition to the primary sex dimorphism genes responsible for anther and carpel development, this enables the identification of secondary sex dimorphisms, such as genes involved in pigmentation, volatile emission, and differences in catkin phenology, which can also inform differences in vegetative emergence and secondary metabolites. By using network analysis and incorporating genomic data through eQTL, we can hypothesize how the SDR may regulate differential gene expression in catkins. Nearly two thirds of all expressed genes in the floral tissue exhibited differential expression between males and females. This number is due in part to the large sample size of 159 individuals, whereby there is enough statistical power to detect even slight differences in expression. Nevertheless, over 21% of the expressed genes showed at least two-fold expression differences between sexes, providing evidence of global expression differences, which would require robust transcriptional regulation, ultimately leading back to the sex determinant genes in the SDR. These genes provide important clues about the regulation of sex determination in this species and the molecular mechanism responsible for dioecy and floral sex dimorphism, as described in more detail below.

### Type A Response Regulator (ARR17)

Four copies of *ARR17*, a type A cytokinin response regulator, in the SDR show high levels of expression in female *S. purpurea*: Sapur.15WG073500, Sapur.15WG073900, Sapur.15WG074000, and Sapur.15WG075200. Two additional copies of *ARR17* are present on chr19 but are not differentially expressed. The cytokinin signaling pathway has been proposed as a common pathway for sex determination in angiosperms^10^. Cytokinin response regulators serve as feminizing factors in *Actinidia*, where they are master regulators^29^, and *Diospyros*, where they act as top-level regulators downstream of the SDR^2,5^. There is recent evidence implicating *ARR17* as the master regulator of sex in the closely related genus *Populus*, where it may function as a feminizing factor whose expression is suppressed in males by small RNAs^4^.

The presence of two complete copies of *ARR17* on *S. purpurea* chr19, expressed in both males and females, suggests that the dosage of *ARR17* (eight copies in females versus four in males) may have a role in sex determination in willow. Interestingly, these findings suggest a different mechanism for *ARR17* than the leading model for sex determination in *Populus* proposed by Müller et al. (2020)^4^. They proposed that functional *ARR17* in *P. alba* (ZW) is a feminizing factor, and in XY species *ARR17* is silenced by inverted repeats on the Y chromosome through the RNA directed DNA methylation (RDDM) pathway, leading to a male phenotype. To confirm this, they silenced the *ARR17* gene in an early-flowering female line and observed male flowers in tissue culture^4^. We found no evidence of an *ARR17* RNA interference mechanism in *S. purpurea* catkins. *Salix purpurea* has a similar truncated inverted repeat of *ARR17* on chr15Z, but we did not observe small RNAs mapping to the *ARR17* genes and their proximal regions, nor to the *ARR17* homologs on *Salix* chr19. There were also no differential methylated regions in the putative promoter regions of any of the *ARR17* genes, and *S. purpurea* males show expression of the *ARR17* copies on chr19, whereas in *P. trichocarpa* males, there is no *ARR17* expression. Furthermore, Carlson et al. (2017) did not find that ARR17 was differentially expressed in shoot tips containing floral primordia, indicating that this mechanism is not present at an earlier floral development stage either^27^. Taken together, these results suggest that the RNA-interference mechanism of *ARR17* may be absent in *S. purpurea*.

The observation of *ARR17* expression in both male (chr19) and female (chr19 and chr15W) *S. purpurea*, combined with a lack of small RNA loci in these same regions, demonstrates that the *Salix* sex determination mechanism is likely different from the model proposed by Müller et al. (2020)^4^. Instead, our data suggest that if *ARR17* is a master regulator in *S. purpurea*, it likely involves a unique mechanism, possibly through gene dosage, such that a threshold of *ARR17* expression must be reached to activate a switch from male to female development. Alternatively, there may be another feature in the *S. purpurea* SDR that is suppressing this silencing mechanism, one such candidate is the adjacent *AGO4* homolog described below.

### ARGONAUTE 4 (AGO4)

A single copy of an Arabidopsis *AGO4* homolog, Sapur.15WG074400, is present within the *ARR17* inverted repeat region of the chr15W SDR^14^ exhibits a log2 M:F expression of −7.94, and has a cis eQTL in the SDR. *AGO4* is a component of the RNA-Induced Silencing Complex (RISC) in the RNA-dependent DNA Methylation (RDDM) pathway, where it binds small RNAs and silences mRNA^30^. In the bisulfite sequencing data, nearly three times as many regions showed increased methylation in males (1465) compared to females (553), supporting that methylation activity is downregulated in females and may have a role in mediating sex dimorphisms (Fig. 2). The SDR *AGO4* gene appears to be truncated to only 79 amino acids in length compared to five other catkin-expressed *AGO4* homologs in *S. purpurea*, which are 893 to 922 amino acids, and has multiple indels and substitutions when aligned (Fig. S4A). The most similar AGO4 paralog to Sapur.15WG074400 by MUSCLE multiple sequence alignment is Sapur.008G00580 which has a nearly seven-fold greater expression in males (Fig. S4B, Table S11). We speculate it is possible that the truncated version of *AGO4* is interfering with expression of the full-length Sapur.008G00580 in males by a long non-coding RNA. This could have wide-ranging effects on sexually dimorphic gene expression and could explain the decreased genome wide methylation observed in females. The findings from the bisulfite - sequencing data indicate that methylation is globally reduced in females, suggesting the possibility that the RDDM pathway may be compromised in females. We hypothesize that the Sapur.15WG074400 could be competing for binding of siRNAs with a full-length *AGO4* and sequestering male-specific RDDM in females. Such a mechanism could also explain *ARR17* expression levels, and why no small RNAs were observed mapping to *ARR17* in *Salix*, despite evidence for this mechanism in *Populus*.

### Double Stranded RNA binding protein 1 (DRB1)

A copy of a *DRB1* homolog, Sapur.15WG074300, is also located in the *ARR17* inverted repeat region of the chr15W SDR and is adjacent to *AGO4*. In Arabidopsis, *DRB1* is involved in RNA - mediated post-transcriptional silencing, working directly with *DCL1* in miRNA processing^31^.

RNA modification, RNA metabolic process, and regulation of transcription were all among the enriched GO terms in the female differentially expressed genes and could be regulated directly or indirectly by *DRB1*.

### Transcription Factor GATA15

A female expressed homolog of *GATA15*, Sapur.15WG062800, is located in the W-specific region of the SDR and shows a *cis*-eQTL association with polymorphisms on Chr15. *GATA15* is a transcriptional regulator that binds GAT or GATA motifs in gene promoters and is involved in cell differentiation, morphogenesis, and development^32^. This is consistent with the GO enrichment analysis of female expressed genes, which contains many significant terms related to morphogenesis and development. Furthermore, chr15 *GATA15* was found by Carlson et al. (2017) to be differentially expressed in F_1_ *S. purpurea* shoot tips containing floral primordia^27^, suggesting that this may be the earliest cue for floral sex differentiation, which would implicate it as a master regulator gene. While functional genomics data is required to elucidate its precise function, its expression in females both during floral differentiation and catkin emergence suggests that it could be directly involved in gynoecium development.

### Genes of Unknown Function

Four genes were identified that fit the criteria for candidate master regulator genes, but whose functions are not known or whose annotations are insufficient for further analysis. These included Sapur.15WG068800, a CCHC-type zinc finger, and three hypothetical proteins: Sapur.15WG075300, Sapur.15WG074900, and Sapur.15WG075700.

While there is mounting evidence pointing towards *ARR17* as the master regulator in *Populus* spp.^4,24,25^ the evidence for different expression profiles of *ARR17* in *Salix*, as well as the presence of additional candidate genes, suggests that the mechanism may be more complicated or altogether different in *Salix*. Nevertheless, expression data from the *ARR17* homologs in *S. purpurea* does support a possible role in sex determination, either as a single gene master regulator or part of a two-gene system in conjunction with another master regulator, and would provide further evidence to support cytokinin response as a common mechanism for dioecy in angiosperms, as suggested by Montalvão et al^10^. Further functional genomics studies will be necessary to elucidate the precise functions of candidate master regulators and their role in sex determination.

### SDR Regulation of Floral Sex Dimorphism

Among the floral development genes with eQTL in the SDR were homologs of *AGL11* and *AGL32* (ovule development and seed formation)^33^, *AGL29* and *AGL30* (pollen development)^34^, *AGL6* (floral meristem differentiation and gamete development)^35^, *TOC1* and *CONSTANS* (flowering response to environmental cues)^36^ (Table S10). While differential expression of MADS-box genes is expected in floral tissues, their association with the SDR through eQTL, even after accounting for sex as a covariate, suggests that the SDR may have a direct role in controlling expression of these genes and the resulting primary sex dimorphisms. These floral development genes may have specific roles as “entry points” for the SDR into the floral development pathway to regulate the development of a particular sex (Fig. 4). Cronk and Müller (2020) proposed that *ARR17* may act as a feminizing master regulator in *Populus* through the suppression of *PISTILLATA* (*PI*) or *APETALLA3* (*AP3*) MADS-box genes^25^. Importantly, there was no association of either *PI* or *AP3* expression with genomic variation in the SDR, despite the fact that *PI* shows very high levels of expression in males (log2 M:F 13.21) (Table S11), further suggesting that the mechanism of sex determination in *Salix* may be different from *Populus*.

**Figure 4.**
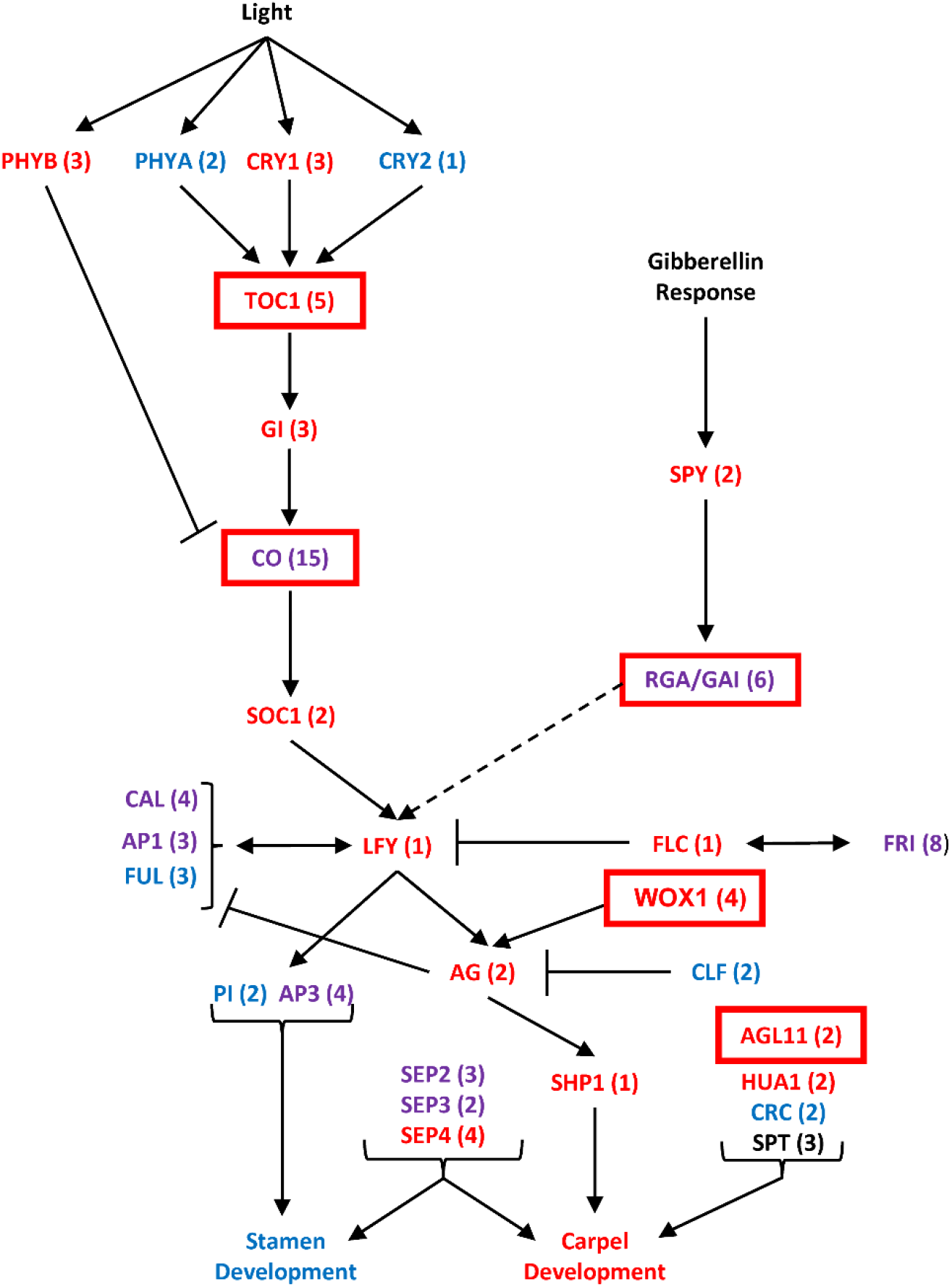
Expression network (adapted from: Blazquez, 2000 and De Bodt et al. 2006) displaying known roles of flowering pathways and genes in Arabidopsis whose homologs are expressed in *S. purpurea* catkins. The number of copies of each gene in *S. purpurea* is shown in parentheses. Female differentially expressed genes are shown in red, male differentially expressed in blue, and genes with both male and female differentially expressed copies in purple. Genes with eQTL associations to the sex determination region are outlined in red boxes and represent possible entry points for SDR regulation of the flowering pathway.

eQTL analysis revealed several loci with an exceptionally high number of *trans*-eQTL (Fig. 2). Intriguingly, The hotspot with the greatest number of *trans*-eQTLs is located in the region on chr07 homologous to the *S. nigra* SDR^*12*^, which could either point towards its role as an ancestral SDR in *Salix*, or explain a fitness advantage of a chr07 SDR in *S. nigra* when linked with sex-dimorphism genes in this region. Approximately 250 kb from this locus on chr07 is Sapur.007G068100, a homolog of *AGL32*, a MADS-box gene involved in ovule development^33^. The expression of this gene is associated in *trans* with the chromosome 15 SDR region, further supporting its role as a top-level regulatory gene under direct regulation by the SDR.

Compounds produced from the phenylpropanoid and terpenoid pathways are well characterized in Salicaceae, and there is evidence that floral volatile, terpenoid, and phenolic glycoside profiles differ substantially between males and females, which affects both pollinator attraction and herbivory, traits that are likely to be evolutionary drivers of dioecy and affect cultivar yield^22,23^. In support of this, we identified five terpenoid pathway genes and 15 phenylpropanoid pathway genes with eQTL in the SDR. These include both genes involved in the core phenylpropanoid pathway, and biosynthesis of specific compounds, including naringenin, flavenol glucosides, and sesquiterpenoids (Table S10). This provides evidence supporting a direct link between the SDR and synthesis of these compounds, by an as-yet unknown mechanism.

### Small RNA Regulation of Sex Dimorphism

Of the 45 identified miRNA loci, 18 were male differentially expressed and 7 were female differentially expressed. Among the putative targets of these miRNAs were dicer-like genes, squamosa promoter-like genes, and auxin response factors, and transcription factors (Table 1) providing evidence that miRNAs are likely to be a component of floral sex dimorphism regulation. Notably, five miRNAs were identified that had no match with any small RNAs in the PmiREN database and could represent genus or species specific micro RNAs. Expression results showed that four miR156 and two miR172 homologs have greater expression in male floral tissue. In Arabidopsis, miR156 and miR172 interact to form a gradient that regulates vegetative to floral meristem transition through the targeting of squamosa promoter-like genes (*SPL*)^28^. While all copies of both miR172 and miR156 show male biased expression in catkins, the overall expression of miR156 is greater than miR172 in males. The majority of *SPL* genes that are targeted are female upregulated in *S. purpurea*, including an *SPL4* homolog on chr07 (Sapur.007G123900) that also shows increased methylation in males in its promoter region. These data support that the miR156/172 pathway is upregulated in male catkins and may be responsible for sex dimorphisms (Table S11). This pathway may play a role in male floral tissue development or differentiation.

Intriguingly, one copy of miR156 is located in the SDR region unique to chr15Z. Alignment of the chr15Z miR156 precursor sequence to chr15W reveals a single indel is responsible for this chr15Z-specific mapping (Figure S5), which may prevent transcription or processing of this small RNA from chr15W. This could indicate a dosage dependent response in males, which have four copies of this mature miR156 homolog, compared to only three in females. Female upregulated miRNAs included miR403, which targets *AGO2*, and miR162, which targets a dicer-like gene. All of these are involved in small RNA signaling and DNA methylation, which is consistent with the enrichment of transcription regulation terms in the female upregulated genes. This may point towards a role of genome-wide DNA methylation or RNA silencing in regulating sex dimorphism, possibly mediated by *AGO4* or *DRB1*.

## Conclusion

Taken together, RNA-Seq, small RNA-Seq, bisulfite sequencing, and eQTL mapping results suggest that the sex determination mechanism in willow is different from the small RNA - mediated, single-gene sex determination pathway proposed in *Populus*, and reveals that many miRNAs, transcription factors, floral development genes, and secondary metabolism genes are involved in primary and secondary sex dimorphism pathways in *Salix purpurea* catkins. A major factor inhibiting functional genomics research in *Salix* is the lack of a functional transformation system, which precludes direct assessment of gene function in transgenic plants. We show here that using a combination of genome-wide differential expression, co-expression, eQTL, small RNA, and genomic variation can provide strong evidence supporting the involvement of these proposed candidate master regulator genes in sex determination.

## Materials and Methods

### Plant material and growing conditions

The F_1_ family (Family 82) was generated from a cross between female *S. purpurea* clone 94006 and male clone 94001, both collected from different naturalized *S. purpurea* populations in Upstate NY (Fig. S6)^37,38^. A female F_1_ individual, *S. purpurea* ‘Wolcott’ (clone 9882-41) and male F_1_ individual, *S. purpurea* ‘Fish Creek’ (clone 9882-34), were crossed to generate the F_2_ *S. purpurea* family (Family 317). All progeny individuals and their parents were planted in nursery beds at Cornell AgriTech, Geneva, NY. For additional information on the F_1_ and F_2_ families and their parents, see Carlson et al. (2019)^39^. Details on phenological stage of catkin collection can be found in the Supplementary Methods.

### Library Preparation and Sequencing

Catkins of 90 males and 90 females from the 317 F_2_ family were collected for RNA-Seq, which, after removing individuals with poor mapping quality, was reduced to 77 females and 82 males. Reads were mapped using STAR^40^ (Table S12), and counts assigned using featureCounts^41^ in R. DESeq2 was used to identify differentially expressed genes^42^. A total of 22 male and 21 female F_2_ progeny, along with the male (94001) and female (94006) grandparents were collected for small RNA sequencing. Mapping and identification of putative miRNAs was done using ShortStacks^43^ and differential expression determined using DESeq2^42^. Six male and six female F_2_ progeny, along with twelve male and twelve female unrelated *S. purpurea* from a diversity panel were collected for bisulfite sequencing. Reads were mapped using Bismark^44^ and differentially methylated regions (DMR) determined using DMRfinder^45^. GBS data were collected on all 317 family F_2_ progeny and used to call variants in TASSEL 5.0^46^. The RNA-Seq mapping and downstream analysis pipelines are summarized in Fig. S7. Full details on library prep, sequencing, and alignment to the reference genome can be found in the Supplementary Methods.

### eQTL Analysis

The R package, MatrixEQTL^47^, was used to map eQTL. Covariates were included for sex, along with the four largest principle components in order to control for underlying data structure and minimize false positive eQTL mapping to the SDR. Each unique SNP/gene association is considered separately as an eQTL, independent of any additional genes that may be associated with that SNP. *Cis*-associations were classified as those SNPs within 1 Mb of their associated gene based on the recommended settings of MatrixEQTL, while *trans*-associations were classified as greater than 1 Mb or on different chromosomes. SNPs within the SDR, spanning from 2.34 Mb to 9.07 Mb on chr15W and 2.34 Mb to 6.70 Mb on chr15Z^14^, were also considered to be *cis* to all other genes within the SDR, regardless of distance, due to low recombination in this region.

### WGCNA Network Analysis

Library normalized FPKM expression values were calculated from the raw count data using the edgeR package^48^. Weighted gene co-expression network analysis (WGCNA)^49^ was used to construct modules of genes with similar expression values. A clustering of FPKM expression values was generated using hclust, which showed that two male samples (10X-317-161 and 10X-317-020) displayed extreme differences in overall expression patterns from all of the other samples (Fig. S8), and so were removed from downstream analyses. A topological overview matrix was created to perform module identification. In order to correlate sex with module expression, sex was coded numerically with values of −1 for females and 1 for males, thus a module r^2^ near one indicates strong correlation with males, and a module r^2^ near negative one indicates strong correlation with females. The minimum module size threshold was set at 30 genes per module. Identified modules were merged based on similar expression levels. A total of 32 unmerged modules were merged into 24 modules based on similar expression patterns.

### Gene Ontology Enrichment and Pathway Analysis

Gene ontology (GO) enrichment at the lowest levels of the GO hierarchy was calculated in the BiNGO app^50^ in Cytoscape using the Arabidopsis homolog gene models. Arabidopsis gene models were used as there is limited data available on gene function in *Salix*, and gene functional annotations of non-model species are often ultimately based on experimental evidence from Arabidopsis. Furthermore, using Arabidopsis gene models provided compatibility with existing databases, such as TAIR. The background reference was created from the list of Arabidopsis homologs for every transcript model in the *S. purpurea* 94006 v5.1 genome found in the gene annotation information file on Phytozome (https://phytozome-next.jgi.doe.gov/info/Spurpurea_v5_1). The Arabidopsis homologs of the genes present in each module or male:female differential expression cutoff (inclusive of all genes above or below cutoff) were used as the query to determine overrepresented and underrepresented parent and child terms for each gene set. Descriptions of each over and under-represented Gene Ontology ID were generated by BinGO based on TAIR descriptions^36^.

## Acknowledgements

Support for this research was provided by grants (DEB-1542486, DEB-1542599) from the National Science Foundation and from the USDA National Institute for Food and Agriculture (2015-67009-23957). The work conducted by the DOE JGI is supported by the Office of Science of the US Department of Energy under Contract no. DE-AC02-05CH11231. Partial support for BLH was provided by a Cornell AgriTech Extension and Outreach Assistantship. The authors would like to thank Matt Christiansen, Curt Carter, Rebecca Wilk, Lauren Carlson, Jane Petzoldt, Dawn Fishback, and Sam Knopka for their technical assistance with sample collection and field trial maintenance and Alex Harkess and Xiohan Yang for critical feedback on early and late drafts of the manuscript, respectively.

## Author Contributions

BLH and CHC contributed equally to the manuscript. BLH designed the data analysis workflow, performed data analysis and interpretation and wrote the manuscript. CHC contributed to experimental design, performed catkin collections, total RNA and small RNA isolations, data collection, and bioinformatics. FEG contributed to experimental design and data collection. GF contributed to development of custom mapping genome for RNA-Seq data. LBS, SPD, MSO and GAT obtained funding and contributed to the experimental design, execution of various stages of the research, and preparation of the manuscript. SPD and MSO made substantial revisions and comments on late versions of the manuscript. AL, AS, KB and LS carried out transcriptome sequencing.

## Data Availability

Raw RNA-Seq, small RNA-Seq, and bisulfite sequencing data are available at the JGI genome portal under proposal 1690, as well as on the NCBI sequence read archive (SRX5027565 - SRX5027793, SRX3886725-SRX3886746, SRX3987232-SRX3987253). Code used in the mapping and analysis are available at https://github.com/Willowpedia.

## Supplementary Material Table Legends

**Supplementary Table S1**. Overrepresented terms among the significant male differentially expressed genes with a value of log_2_ >1.

**Supplementary Table S2**. Underrepresented terms among the significant male differentially expressed genes with a value of log2 >1.

**Supplementary Table S3**. Overrepresented terms among the significant female differentially expressed genes with a value of log2 <-1

**Supplementary Table S4**. Underrepresented terms among the significant female differentially expressed genes with a value of log2<-1

**Supplementary Table S5**. Summary of the four miR156 loci, including the sequence of the most abundant RNA at each locus and the number of alignments for each small RNA length.

**Supplementary Table S6**. All differentially methylated regions (DMR) among 24 diverse S. purpurea and 12 F_2_ individuals.

**Supplementary Table S7**. All 170 genes with DMRs in their putative promoter regions

**Supplementary Table S8**. Overrepresented terms in the female correlated “purple” module.

**Supplementary Table S9**. Overrepresented terms in the male correlated “cyan” module.

**Supplementary Table S10**. All genes associated with the SDR through eQTL analysis with annotations involving transcriptional regulation, floral development, or secondary metabolism, or RNA splicing and regulation, ordered by Gene ID number. Dominance deviations from the additive model are shown, along with their significance.

**Supplementary Table S11**. All differential expression analysis and WGCNA module membership for each gene, along with annotation info obtained from Phytozome 13.

**Supplementary Table S12**. Read mapping statistics for all RNA-Seq libraries. Note that some samples had two library preps.

## Figure Legends

**Supplementary Figure S1**. Principal component analysis plot of RNA-Seq data from catkins of 180 F2 S. purpurea family progeny, prior to removing samples with low mapping rate (<70%). Males are coded in blue and females in magenta.

**Supplementary Figure S2**. Read length distribution of small RNA raw reads after QC.

**Supplementary Figure S3**. MatrixeQTL results from each of the 136 genes identified with SDR eQTL that have transcription, floral development, secondary metabolism, or RNA regulation annotations. “S15” refers to loci on the chr15Z SDR region, while “S20” refers to loci on the chr15W SDR. Males are coded in blue and females in magenta

**Supplementary Figure S4**. (**A**) MUSCLE Multiple Sequence Alignment of the six floral expressed *AGO4* protein sequences, (**B**) Neighbor-joining phylogenetic tree of MSA alignment without distance corrections produced by MUSCLE

**Supplementary Figure S5**. (**A**) Structural alignment of Cluster 187938, a miR156 homolog on chr15Z, for each aligned sequence “l” indicates the length, and “a” the number of reads aligning with that particular sequence. (**B**) BLAST alignment of the chr15Z miR156 precursor sequence to chr15W shows a single indel that results on chr15Z specific mapping.

**Supplementary Figure S6**. Pedigree of Family ‘317’ used for RNA-Seq and smRNA analysis.

**Supplementary Figure S7**. Diagram summarizing the RNA-Seq analysis pipeline and workflow, including RNA-Seq and GBS read alignments, eQTL mapping, differential expression analysis, and network analysis.

**Supplementary Figure S8**. Clustering of samples analyzed using WGCNA. 10X-317-161 and 10X-317-020 were clear outliers and were removed from further network analysis.

