## Supplementary Tables S1-S4, S8-S9, S12; Supplementary Figs. S1-S2, S4-S8 for "Integrative genomics reveals paths to sex dimorphism in *Salix purpurea* L."

**Supplementary Table S1.** Overrepresented terms among the significant male differentially expressed genes with a value of log$_{2}$>1.

**Supplementary Table S2.** Underrepresented terms among the significant male differentially expressed genes with a value of log2 >1.

**Supplementary Table S3.** Overrepresented terms among the significant female differentially expressed genes with a value of log2<-1

**Supplementary Table S4.** Underrepresented terms among the significant female differentially expressed genes with a value of log2<-1

**Supplementary Table S8.** Overrepresented terms in the female correlated “purple” module.

**Supplementary Table S9.** Overrepresented terms in the male correlated “cyan” module.

**Supplementary Table S12**. Read mapping statistics for all RNA-Seq libraries. Note that some samples had two library preps.

**Figure Legends**

**Supplementary Figure S1.** Principal component analysis plot of RNA-Seq data from catkins of 180 F2 S. purpurea family progeny, prior to removing samples with low mapping rate (<70%). Males are coded in blue and females in magenta.

**Supplementary Figure S2.** Read length distribution of small RNA raw reads after QC.

**Supplementary Figure S4.** (**A**) MUSCLE Multiple Sequence Alignment of the six floral expressed *AGO4* protein sequences, (**B**) Neighbor-joining phylogenetic tree of MSA alignment without distance corrections produced by MUSCLE

**Supplementary Figure S5.** (**A**) Structural alignment of Cluster 187938, a miR156 homolog on chr15Z, for each aligned sequence “l” indicates the length, and “a” the number of reads aligning with that particular sequence. (**B**) BLAST alignment of the chr15Z miR156 precursor sequence to chr15W shows a single indel that results on chr15Z specific mapping.

**Supplementary Figure S6.** Pedigree of Family ‘317’ used for RNA-Seq and smRNA analysis.

**Supplementary Figure S7.** Diagram summarizing the RNA-Seq analysis pipeline and workflow, including RNA-Seq and GBS read alignments, eQTL mapping, differential expression analysis, and network analysis.

**Supplementary Figure S8.** Clustering of samples analyzed using WGCNA. 10X-317-161 and 10X-317-020 were clear outliers and were removed from further network analysis.

### Supplementary Table S1

| GO ID | GO Description | Genes Present | FDR |
|---|---|---|---|
| 272 | polysaccharide catabolic process | 28 | 7.25E-04 |
| 5975 | carbohydrate metabolic process | 131 | 4.29E-06 |
| 5976 | polysaccharide metabolic process | 52 | 1.09E-03 |
| 6468 | protein amino acid phosphorylation | 95 | 7.82E-04 |
| 6629 | lipid metabolic process | 92 | 9.42E-03 |
| 6793 | phosphorus metabolic process | 135 | 2.39E-03 |
| 6796 | phosphate metabolic process | 135 | 2.39E-03 |
| 6811 | ion transport | 54 | 7.25E-04 |
| 6812 | cation transport | 47 | 1.13E-04 |
| 6813 | potassium ion transport | 12 | 1.03E-02 |
| 6816 | calcium ion transport | 10 | 2.48E-02 |
| 6873 | cellular ion homeostasis | 23 | 1.03E-02 |
| 6885 | regulation of pH | 13 | 7.42E-04 |
| 6970 | response to osmotic stress | 27 | 1.81E-02 |
| 7276 | gamete generation | 6 | 3.73E-02 |
| 9269 | response to desiccation | 7 | 1.36E-02 |
| 9414 | response to water deprivation | 20 | 4.73E-02 |
| 9415 | response to water | 20 | 4.73E-02 |
| 9555 | pollen development | 20 | 6.94E-04 |
| 9651 | response to salt stress | 25 | 1.58E-02 |
| 9690 | cytokinin metabolic process | 7 | 3.81E-02 |
| 10208 | pollen wall assembly | 7 | 2.00E-02 |
| 10584 | pollen exine formation | 6 | 3.73E-02 |
| 10927 | cellular component assembly involved in morphogenesis | 7 | 2.00E-02 |
| 15672 | monovalent inorganic cation transport | 16 | 2.22E-02 |
| 15674 | di-, tri-valent inorganic cation transport | 19 | 1.30E-02 |
| 16042 | lipid catabolic process | 32 | 3.04E-03 |
| 16052 | carbohydrate catabolic process | 46 | 2.15E-04 |
| 16103 | diterpenoid catabolic process | 4 | 1.03E-02 |
| 16139 | glycoside catabolic process | 7 | 9.42E-03 |
| 16145 | S-glycoside catabolic process | 4 | 2.71E-02 |
| 16310 | phosphorylation | 114 | 2.00E-02 |
| 19748 | secondary metabolic process | 36 | 1.81E-02 |
| 19759 | glycosinolate catabolic process | 4 | 2.71E-02 |
| 19762 | glucosinolate catabolic process | 4 | 2.71E-02 |
| 19953 | sexual reproduction | 13 | 9.42E-03 |
| 30001 | metal ion transport | 46 | 7.12E-06 |
| 30198 | extracellular matrix organization | 7 | 2.71E-02 |

| | | | |
|---|---|---|---|
| 32504 | multicellular organism reproduction | 10 | 2.78E-02 |
| 33499 | galactose catabolic process via UDP-galactose | 5 | 4.73E-02 |
| 42221 | response to chemical stimulus | 159 | 1.58E-02 |
| 42542 | response to hydrogen peroxide | 16 | 3.73E-02 |
| 42545 | cell wall modification | 22 | 6.81E-04 |
| 42592 | homeostatic process | 51 | 9.87E-03 |
| 42743 | hydrogen peroxide metabolic process | 14 | 4.19E-02 |
| 42744 | hydrogen peroxide catabolic process | 14 | 4.19E-02 |
| 43062 | extracellular structure organization | 7 | 2.71E-02 |
| 44247 | cellular polysaccharide catabolic process | 18 | 1.17E-02 |
| 44262 | cellular carbohydrate metabolic process | 87 | 1.51E-02 |
| 44264 | cellular polysaccharide metabolic process | 41 | 1.78E-02 |
| 44273 | sulfur compound catabolic process | 5 | 4.73E-02 |
| 44275 | cellular carbohydrate catabolic process | 36 | 3.04E-03 |
| 45229 | external encapsulating structure organization | 9 | 2.20E-03 |
| 45487 | gibberellin catabolic process | 4 | 1.03E-02 |
| 45488 | pectin metabolic process | 22 | 1.49E-03 |
| 45490 | pectin catabolic process | 18 | 5.89E-04 |
| 45493 | xylan catabolic process | 5 | 2.78E-02 |
| 48229 | gametophyte development | 25 | 2.61E-03 |
| 48232 | male gamete generation | 6 | 2.50E-02 |
| 48235 | pollen sperm cell differentiation | 6 | 2.50E-02 |
| 48609 | reproductive process in a multicellular organism | 10 | 2.78E-02 |
| 48610 | reproductive cellular process | 18 | 2.39E-03 |
| 48878 | chemical homeostasis | 44 | 8.22E-06 |
| 50801 | ion homeostasis | 40 | 4.29E-06 |
| 51017 | actin filament bundle assembly | 5 | 4.73E-02 |
| 55046 | microgametogenesis | 6 | 2.50E-02 |
| 55066 | di-, tri-valent inorganic cation homeostasis | 22 | 2.50E-02 |
| 55067 | monovalent inorganic cation homeostasis | 14 | 1.06E-03 |
| 55080 | cation homeostasis | 36 | 3.69E-05 |
| 55082 | cellular chemical homeostasis | 23 | 1.30E-02 |
| 65008 | regulation of biological quality | 84 | 1.30E-02 |
| 70301 | cellular response to hydrogen peroxide | 14 | 4.73E-02 |
| 70838 | divalent metal ion transport | 12 | 3.09E-02 |
| 71554 | cell wall organization or biogenesis | 80 | 2.56E-10 |
| 71555 | cell wall organization | 66 | 6.44E-08 |
| 71669 | plant-type cell wall organization or biogenesis | 21 | 4.50E-02 |

### Supplementary Table S2

| GO ID | GO Description | Genes Present | FDR |
|---|---|---|---|
| 375 | RNA splicing, via transesterification reactions | 1 | 3.64E-02 |
| 377 | RNA splicing, via transesterification reactions with bulged adenosine as nucleophile | 1 | 3.64E-02 |
| 6139 | nucleobase, nucleoside, nucleotide and nucleic acid metabolic process | 43 | 6.81E-25 |
| 6259 | DNA metabolic process | 11 | 9.50E-05 |
| 6260 | DNA replication | 2 | 9.57E-03 |
| 6261 | DNA-dependent DNA replication | 0 | 3.21E-03 |
| 6281 | DNA repair | 6 | 6.16E-03 |
| 6364 | rRNA processing | 0 | 1.66E-03 |
| 6396 | RNA processing | 9 | 9.19E-11 |
| 6397 | mRNA processing | 6 | 4.94E-05 |
| 6399 | tRNA metabolic process | 2 | 3.21E-02 |
| 6403 | RNA localization | 0 | 2.63E-02 |
| 6412 | translation | 11 | 1.56E-05 |
| 6511 | ubiquitin-dependent protein catabolic process | 11 | 4.35E-03 |
| 6605 | protein targeting | 3 | 2.15E-04 |
| 6807 | nitrogen compound metabolic process | 73 | 1.07E-24 |
| 6913 | nucleocytoplasmic transport | 0 | 1.51E-02 |
| 6974 | response to DNA damage stimulus | 7 | 4.35E-03 |
| 6996 | organelle organization | 55 | 2.87E-03 |
| 7005 | mitochondrion organization | 2 | 7.99E-03 |
| 8152 | metabolic process | 596 | 4.43E-02 |
| 8380 | RNA splicing | 1 | 4.49E-04 |
| 9058 | biosynthetic process | 149 | 9.42E-03 |
| 9059 | macromolecule biosynthetic process | 53 | 4.17E-05 |
| 9108 | coenzyme biosynthetic process | 1 | 4.33E-02 |
| 9451 | RNA modification | 4 | 5.90E-06 |
| 10467 | gene expression | 27 | 3.95E-19 |
| 16070 | RNA metabolic process | 19 | 4.04E-19 |
| 16071 | mRNA metabolic process | 7 | 5.90E-06 |
| 16072 | rRNA metabolic process | 0 | 5.94E-04 |
| 17038 | protein import | 0 | 4.76E-04 |
| 18130 | heterocycle biosynthetic process | 1 | 2.38E-03 |
| 19941 | modification-dependent protein catabolic process | 13 | 1.55E-02 |
| 22613 | ribonucleoprotein complex biogenesis | 2 | 5.07E-07 |
| 22618 | ribonucleoprotein complex assembly | 2 | 4.54E-02 |
| 30163 | protein catabolic process | 22 | 2.49E-02 |
| 33365 | protein localization in organelle | 0 | 1.08E-04 |

| | | | |
|---|---|---|---|
| 34470 | ncRNA processing | 3 | 1.73E-03 |
| 34613 | cellular protein localization | 22 | 4.43E-02 |
| 34621 | cellular macromolecular complex subunit organization | 11 | 9.09E-05 |
| 34622 | cellular macromolecular complex assembly | 10 | 1.36E-03 |
| 34641 | cellular nitrogen compound metabolic process | 62 | 2.53E-27 |
| 34645 | cellular macromolecule biosynthetic process | 52 | 2.49E-05 |
| 34660 | ncRNA metabolic process | 2 | 5.31E-06 |
| 42254 | ribosome biogenesis | 1 | 5.22E-06 |
| 42440 | pigment metabolic process | 1 | 4.33E-02 |
| 43170 | macromolecule metabolic process | 290 | 2.67E-08 |
| 43632 | modification-dependent macromolecule catabolic process | 13 | 1.00E-02 |
| 43933 | macromolecular complex subunit organization | 15 | 2.97E-04 |
| 44237 | cellular metabolic process | 451 | 3.55E-07 |
| 44238 | primary metabolic process | 481 | 4.43E-02 |
| 44249 | cellular biosynthetic process | 137 | 1.36E-03 |
| 44257 | cellular protein catabolic process | 18 | 1.76E-02 |
| 44260 | cellular macromolecule metabolic process | 262 | 3.36E-09 |
| 44265 | cellular macromolecule catabolic process | 23 | 6.53E-03 |
| 44271 | cellular nitrogen compound biosynthetic process | 19 | 1.20E-02 |
| 46483 | heterocycle metabolic process | 15 | 4.54E-02 |
| 46907 | intracellular transport | 25 | 2.81E-04 |
| 50657 | nucleic acid transport | 0 | 4.33E-02 |
| 50658 | RNA transport | 0 | 4.33E-02 |
| 51169 | nuclear transport | 0 | 1.51E-02 |
| 51186 | cofactor metabolic process | 13 | 5.81E-03 |
| 51188 | cofactor biosynthetic process | 1 | 1.00E-04 |
| 51236 | establishment of RNA localization | 0 | 4.33E-02 |
| 51276 | chromosome organization | 10 | 1.24E-02 |
| 51603 | proteolysis involved in cellular protein catabolic process | 18 | 2.14E-02 |
| 51641 | cellular localization | 37 | 6.53E-03 |
| 51649 | establishment of localization in cell | 33 | 6.54E-03 |
| 65003 | macromolecular complex assembly | 14 | 3.61E-03 |
| 70727 | cellular macromolecule localization | 22 | 3.50E-02 |
| 90304 | nucleic acid metabolic process | 29 | 6.81E-25 |

### Supplementary Table S3

| GO ID | GO Description | Genes Represented | FDR |
|---|---|---|---|
| 3002 | regionalization | 12 | 2.56E-03 |
| 6355 | regulation of transcription, DNA-dependent | 109 | 2.74E-04 |
| 7275 | multicellular organismal development | 92 | 1.34E-02 |
| 7389 | pattern specification process | 13 | 6.35E-03 |
| 9719 | response to endogenous stimulus | 58 | 3.84E-02 |
| 9755 | hormone-mediated signaling pathway | 42 | 4.61E-02 |
| 9889 | regulation of biosynthetic process | 116 | 4.86E-04 |
| 9943 | adaxial/abaxial axis specification | 4 | 2.61E-02 |
| 9944 | polarity specification of adaxial/abaxial axis | 4 | 2.61E-02 |
| 9955 | adaxial/abaxial pattern formation | 4 | 2.61E-02 |
| 10103 | stomatal complex morphogenesis | 5 | 4.46E-02 |
| 10374 | stomatal complex development | 8 | 1.91E-03 |
| 10468 | regulation of gene expression | 111 | 9.35E-03 |
| 10556 | regulation of macromolecule biosynthetic process | 113 | 7.23E-04 |
| 19219 | regulation of nucleobase, nucleoside, nucleotide and nucleic acid metabolic process | 113 | 4.86E-04 |
| 19222 | regulation of metabolic process | 127 | 4.69E-03 |
| 22621 | shoot system development | 24 | 7.68E-06 |
| 30154 | cell differentiation | 35 | 5.34E-05 |
| 31323 | regulation of cellular metabolic process | 124 | 2.37E-05 |
| 31326 | regulation of cellular biosynthetic process | 115 | 3.35E-06 |
| 32501 | multicellular organismal process | 103 | 1.47E-05 |
| 32502 | developmental process | 110 | 1.04E-05 |
| 45449 | regulation of transcription | 109 | 4.26E-07 |
| 48367 | shoot development | 24 | 9.90E-04 |
| 48513 | organ development | 39 | 1.76E-03 |
| 48731 | system development | 40 | 1.04E-03 |
| 48869 | cellular developmental process | 43 | 4.62E-03 |
| 51171 | regulation of nitrogen compound metabolic process | 114 | 4.86E-04 |
| 51252 | regulation of RNA metabolic process | 109 | 4.86E-04 |
| 60255 | regulation of macromolecule metabolic process | 115 | 1.68E-02 |
| 65001 | specification of axis polarity | 4 | 2.61E-02 |
| 80090 | regulation of primary metabolic process | 115 | 4.62E-03 |

#### Supplementary Table S4

| GO ID | GO Description | Genes Present | FDR |
|---|---|---|---|
| 5996 | monosaccharide metabolic process | 2 | 3.45E-02 |
| 6139 | nucleobase, nucleoside, nucleotide and nucleic acid metabolic process | 29 | 1.22E-09 |
| 6396 | RNA processing | 5 | 2.98E-05 |
| 6412 | translation | 5 | 2.47E-03 |
| 6511 | ubiquitin-dependent protein catabolic process | 2 | 1.36E-03 |
| 6732 | coenzyme metabolic process | 2 | 1.86E-02 |
| 6807 | nitrogen compound metabolic process | 52 | 5.93E-08 |
| 6810 | transport | 51 | 1.85E-02 |
| 6996 | organelle organization | 26 | 1.05E-02 |
| 8104 | protein localization | 12 | 2.31E-03 |
| 8152 | metabolic process | 296 | 9.87E-05 |
| 9058 | biosynthetic process | 72 | 5.33E-03 |
| 9059 | macromolecule biosynthetic process | 28 | 5.44E-03 |
| 9451 | RNA modification | 1 | 8.80E-04 |
| 9987 | cellular process | 382 | 1.74E-04 |
| 10467 | gene expression | 17 | 2.21E-08 |
| 15031 | protein transport | 10 | 1.57E-03 |
| 16070 | RNA metabolic process | 14 | 1.69E-07 |
| 16071 | mRNA metabolic process | 5 | 2.13E-02 |
| 16192 | vesicle-mediated transport | 5 | 2.66E-02 |
| 19941 | modification-dependent protein catabolic process | 2 | 1.16E-03 |
| 22613 | ribonucleoprotein complex biogenesis | 1 | 1.64E-03 |
| 33036 | macromolecule localization | 20 | 2.32E-02 |
| 34641 | cellular nitrogen compound metabolic process | 46 | 7.27E-09 |
| 34645 | cellular macromolecule biosynthetic process | 28 | 5.68E-03 |
| 34660 | ncRNA metabolic process | 0 | 6.88E-04 |
| 42254 | ribosome biogenesis | 0 | 1.75E-03 |
| 43170 | macromolecule metabolic process | 153 | 1.36E-05 |
| 43412 | macromolecule modification | 72 | 2.48E-02 |
| 43632 | modification-dependent macromolecule catabolic process | 2 | 8.38E-04 |
| 44237 | cellular metabolic process | 212 | 1.84E-10 |
| 44238 | primary metabolic process | 232 | 6.39E-05 |
| 44249 | cellular biosynthetic process | 67 | 2.31E-03 |
| 44257 | cellular protein catabolic process | 5 | 4.37E-03 |
| 44260 | cellular macromolecule metabolic process | 124 | 2.98E-09 |
| 44262 | cellular carbohydrate metabolic process | 16 | 2.26E-02 |
| 44265 | cellular macromolecule catabolic process | 5 | 1.42E-04 |
| 44267 | cellular protein metabolic process | 91 | 5.96E-03 |

| | | | |
|---|---|---|---|
| 45184 | establishment of protein localization | 10 | 1.36E-03 |
| 46907 | intracellular transport | 12 | 8.40E-03 |
| 51179 | localization | 54 | 1.74E-02 |
| 51186 | cofactor metabolic process | 4 | 7.41E-03 |
| 51234 | establishment of localization | 52 | 1.87E-02 |
| 51603 | proteolysis involved in cellular protein catabolic process | 5 | 4.98E-03 |
| 51641 | cellular localization | 13 | 1.21E-03 |
| 51649 | establishment of localization in cell | 13 | 5.44E-03 |
| 90304 | nucleic acid metabolic process | 24 | 2.83E-08 |

### Supplementary Table S8

| GO ID | GO Description | Genes Present | FDR |
|---|---|---|---|
| 6139 | nucleobase, nucleoside, nucleotide and nucleic acid metabolic process | 472 | 2.71E-05 |
| 6260 | DNA replication | 53 | 2.95E-02 |
| 6261 | DNA-dependent DNA replication | 41 | 4.73E-02 |
| 6334 | nucleosome assembly | 13 | 3.55E-02 |
| 6355 | regulation of transcription, DNA-dependent | 377 | 2.95E-02 |
| 6807 | nitrogen compound metabolic process | 584 | 1.99E-03 |
| 7018 | microtubule-based movement | 27 | 2.95E-02 |
| 9451 | RNA modification | 134 | 3.22E-16 |
| 9698 | phenylpropanoid metabolic process | 36 | 3.73E-02 |
| 16070 | RNA metabolic process | 297 | 4.27E-04 |
| 19219 | regulation of nucleobase, nucleoside, nucleotide and nucleic acid metabolic process | 407 | 2.95E-02 |
| 19685 | photosynthesis, dark reaction | 11 | 2.95E-02 |
| 34641 | cellular nitrogen compound metabolic process | 566 | 1.19E-03 |
| 45449 | regulation of transcription | 378 | 2.95E-02 |
| 51171 | regulation of nitrogen compound metabolic process | 409 | 2.95E-02 |
| 51252 | regulation of RNA metabolic process | 388 | 3.55E-02 |
| 90304 | nucleic acid metabolic process | 409 | 3.84E-06 |

### Supplementary Table S9

| GO ID | GO Description | Genes Present | FDR |
|---|---|---|---|
| 3 | reproduction | 238 | 2.33E-04 |
| 902 | cell morphogenesis | 79 | 3.75E-05 |
| 904 | cell morphogenesis involved in differentiation | 42 | 1.11E-03 |
| 3006 | reproductive developmental process | 205 | 4.73E-03 |
| 5975 | carbohydrate metabolic process | 200 | 9.61E-04 |
| 5996 | monosaccharide metabolic process | 38 | 1.16E-02 |
| 6066 | alcohol metabolic process | 60 | 6.82E-03 |
| 6082 | organic acid metabolic process | 183 | 3.15E-07 |
| 6090 | pyruvate metabolic process | 9 | 1.23E-02 |
| 6091 | generation of precursor metabolites and energy | 75 | 3.15E-07 |
| 6457 | protein folding | 52 | 4.16E-02 |
| 6464 | protein modification process | 299 | 1.73E-03 |
| 6468 | protein amino acid phosphorylation | 202 | 2.03E-02 |
| 6470 | protein amino acid dephosphorylation | 19 | 4.22E-02 |
| 6487 | protein amino acid N-linked glycosylation | 10 | 4.24E-02 |
| 6629 | lipid metabolic process | 178 | 1.95E-08 |
| 6631 | fatty acid metabolic process | 62 | 2.17E-05 |
| 6633 | fatty acid biosynthetic process | 37 | 3.33E-03 |
| 6638 | neutral lipid metabolic process | 6 | 1.85E-02 |
| 6639 | acylglycerol metabolic process | 6 | 1.85E-02 |
| 6641 | triglyceride metabolic process | 6 | 1.85E-02 |
| 6694 | steroid biosynthetic process | 15 | 4.50E-02 |
| 6720 | isoprenoid metabolic process | 39 | 2.60E-02 |
| 6753 | nucleoside phosphate metabolic process | 44 | 2.26E-02 |
| 6775 | fat-soluble vitamin metabolic process | 8 | 1.85E-02 |
| 6778 | porphyrin metabolic process | 26 | 7.76E-04 |
| 6779 | porphyrin biosynthetic process | 21 | 7.76E-04 |
| 6793 | phosphorus metabolic process | 249 | 2.33E-04 |
| 6796 | phosphate metabolic process | 249 | 2.26E-04 |
| 6810 | transport | 439 | 1.63E-16 |
| 6811 | ion transport | 108 | 1.38E-05 |
| 6812 | cation transport | 81 | 1.10E-03 |
| 6820 | anion transport | 23 | 4.24E-02 |
| 6886 | intracellular protein transport | 77 | 3.70E-05 |
| 6888 | ER to Golgi vesicle-mediated transport | 7 | 4.22E-02 |
| 6897 | endocytosis | 7 | 2.57E-02 |

| | | | |
|---|---|---|---|
| 6944 | cellular membrane fusion | 13 | 4.41E-02 |
| 6950 | response to stress | 456 | 5.24E-06 |
| 6970 | response to osmotic stress | 114 | 1.30E-04 |
| 6979 | response to oxidative stress | 68 | 2.61E-02 |
| 7275 | multicellular organismal development | 423 | 1.82E-07 |
| 7398 | ectoderm development | 38 | 3.72E-02 |
| 7568 | aging | 30 | 5.96E-03 |
| 8104 | protein localization | 104 | 5.59E-05 |
| 8152 | metabolic process | 1455 | 2.34E-03 |
| 8202 | steroid metabolic process | 21 | 4.50E-02 |
| 8300 | isoprenoid catabolic process | 7 | 2.57E-02 |
| 8361 | regulation of cell size | 83 | 4.04E-05 |
| 8544 | epidermis development | 38 | 3.72E-02 |
| 8610 | lipid biosynthetic process | 102 | 1.38E-05 |
| 9056 | catabolic process | 172 | 1.10E-03 |
| 9117 | nucleotide metabolic process | 44 | 2.26E-02 |
| 9144 | purine nucleoside triphosphate metabolic process | 21 | 4.50E-02 |
| 9145 | purine nucleoside triphosphate biosynthetic process | 21 | 4.50E-02 |
| 9165 | nucleotide biosynthetic process | 31 | 4.55E-02 |
| 9199 | ribonucleoside triphosphate metabolic process | 21 | 4.50E-02 |
| 9201 | ribonucleoside triphosphate biosynthetic process | 21 | 4.50E-02 |
| 9205 | purine ribonucleoside triphosphate metabolic process | 21 | 4.50E-02 |
| 9206 | purine ribonucleoside triphosphate biosynthetic process | 21 | 4.50E-02 |
| 9266 | response to temperature stimulus | 99 | 4.39E-03 |
| 9314 | response to radiation | 118 | 3.53E-02 |
| 9409 | response to cold | 68 | 1.62E-02 |
| 9414 | response to water deprivation | 58 | 4.20E-03 |
| 9415 | response to water | 58 | 1.13E-02 |
| 9416 | response to light stimulus | 115 | 2.95E-02 |
| 9553 | embryo sac development | 29 | 2.10E-02 |
| 9555 | pollen development | 48 | 4.64E-03 |
| 9606 | tropism | 19 | 1.98E-02 |
| 9607 | response to biotic stimulus | 160 | 6.22E-06 |
| 9617 | response to bacterium | 73 | 1.54E-03 |
| 9628 | response to abiotic stimulus | 327 | 1.16E-09 |
| 9629 | response to gravity | 19 | 6.73E-03 |
| 9630 | gravitropism | 16 | 2.11E-02 |
| 9651 | response to salt stress | 106 | 2.31E-04 |
| 9653 | anatomical structure morphogenesis | 146 | 6.71E-05 |
| 9657 | plastid organization | 35 | 7.65E-03 |

| | | | |
|---|---|---|---|
| 9694 | jasmonic acid metabolic process | 11 | 2.93E-02 |
| 9743 | response to carbohydrate stimulus | 54 | 8.09E-03 |
| 9744 | response to sucrose stimulus | 16 | 8.53E-03 |
| 9756 | carbohydrate mediated signaling | 13 | 2.60E-02 |
| 9767 | photosynthetic electron transport chain | 15 | 3.66E-04 |
| 9773 | photosynthetic electron transport in photosystem I | 9 | 1.23E-02 |
| 9814 | defense response, incompatible interaction | 37 | 3.33E-03 |
| 9817 | defense response to fungus, incompatible interaction | 15 | 2.88E-02 |
| 9826 | unidimensional cell growth | 63 | 4.73E-05 |
| 9832 | plant-type cell wall biogenesis | 23 | 1.18E-02 |
| 9856 | pollination | 50 | 3.14E-04 |
| 9860 | pollen tube growth | 27 | 2.09E-03 |
| 9867 | jasmonic acid mediated signaling pathway | 16 | 2.60E-02 |
| 9887 | organ morphogenesis | 51 | 3.52E-02 |
| 9888 | tissue development | 69 | 4.98E-02 |
| 9913 | epidermal cell differentiation | 38 | 2.90E-02 |
| 9914 | hormone transport | 19 | 3.57E-02 |
| 9926 | auxin polar transport | 18 | 4.05E-02 |
| 9932 | cell tip growth | 33 | 1.03E-03 |
| 9966 | regulation of signal transduction | 19 | 4.22E-02 |
| 9987 | cellular process | 1637 | 5.10E-08 |
| 10033 | response to organic substance | 248 | 8.65E-03 |
| 10035 | response to inorganic substance | 128 | 3.75E-05 |
| 10038 | response to metal ion | 104 | 1.92E-04 |
| 10149 | senescence | 18 | 7.26E-03 |
| 10150 | leaf senescence | 13 | 2.04E-02 |
| 10182 | sugar mediated signaling pathway | 13 | 2.60E-02 |
| 10189 | vitamin E biosynthetic process | 5 | 4.63E-02 |
| 10193 | response to ozone | 11 | 2.93E-02 |
| 10236 | plastoquinone biosynthetic process | 4 | 2.24E-02 |
| 10260 | organ senescence | 14 | 9.48E-03 |
| 10324 | membrane invagination | 7 | 2.57E-02 |
| 10565 | regulation of cellular ketone metabolic process | 12 | 1.48E-02 |
| 10817 | regulation of hormone levels | 39 | 2.10E-02 |
| 15031 | protein transport | 103 | 1.38E-05 |
| 15698 | inorganic anion transport | 18 | 4.56E-02 |
| 15849 | organic acid transport | 25 | 3.96E-02 |
| 15979 | photosynthesis | 55 | 1.87E-09 |
| 15994 | chlorophyll metabolic process | 18 | 3.82E-03 |
| 15995 | chlorophyll biosynthetic process | 15 | 6.48E-04 |

| | | | |
|---|---|---|---|
| 16042 | lipid catabolic process | 25 | 2.00E-03 |
| 16043 | cellular component organization | 242 | 9.51E-05 |
| 16044 | cellular membrane organization | 32 | 3.14E-04 |
| 16049 | cell growth | 82 | 8.30E-06 |
| 16051 | carbohydrate biosynthetic process | 64 | 2.11E-02 |
| 16053 | organic acid biosynthetic process | 88 | 2.42E-03 |
| 16054 | organic acid catabolic process | 26 | 9.98E-03 |
| 16103 | diterpenoid catabolic process | 5 | 2.24E-02 |
| 16115 | terpenoid catabolic process | 7 | 2.57E-02 |
| 16192 | vesicle-mediated transport | 76 | 1.67E-08 |
| 16310 | phosphorylation | 229 | 8.58E-04 |
| 18130 | heterocycle biosynthetic process | 42 | 1.65E-03 |
| 19318 | hexose metabolic process | 29 | 4.53E-02 |
| 19684 | photosynthesis, light reaction | 28 | 2.55E-04 |
| 19685 | photosynthesis, dark reaction | 5 | 2.24E-02 |
| 19725 | cellular homeostasis | 45 | 2.77E-02 |
| 19752 | carboxylic acid metabolic process | 183 | 2.88E-07 |
| 21700 | developmental maturation | 20 | 2.16E-02 |
| 22414 | reproductive process | 233 | 2.73E-04 |
| 22900 | electron transport chain | 28 | 2.31E-05 |
| 23033 | signaling pathway | 150 | 4.60E-02 |
| 23051 | regulation of signaling process | 19 | 4.22E-02 |
| 30003 | cellular cation homeostasis | 23 | 3.71E-02 |
| 30004 | cellular monovalent inorganic cation homeostasis | 6 | 6.81E-03 |
| 30007 | cellular potassium ion homeostasis | 5 | 6.02E-03 |
| 30154 | cell differentiation | 92 | 9.74E-05 |
| 31407 | oxylipin metabolic process | 13 | 2.60E-02 |
| 31408 | oxylipin biosynthetic process | 11 | 4.16E-02 |
| 32501 | multicellular organismal process | 450 | 1.14E-08 |
| 32502 | developmental process | 469 | 1.25E-08 |
| 32535 | regulation of cellular component size | 83 | 4.64E-05 |
| 32787 | monocarboxylic acid metabolic process | 105 | 1.14E-08 |
| 32940 | secretion by cell | 17 | 4.41E-02 |
| 32989 | cellular component morphogenesis | 83 | 9.51E-05 |
| 33013 | tetrapyrrole metabolic process | 27 | 5.18E-04 |
| 33014 | tetrapyrrole biosynthetic process | 24 | 1.08E-04 |
| 33036 | macromolecule localization | 139 | 1.69E-03 |
| 33238 | regulation of cellular amine metabolic process | 5 | 2.24E-02 |
| 34285 | response to disaccharide stimulus | 16 | 1.17E-02 |
| 34613 | cellular protein localization | 77 | 1.57E-04 |

| | | | |
|---|---|---|---|
| 35295 | tube development | 33 | 4.76E-03 |
| 35466 | regulation of signaling pathway | 31 | 2.18E-02 |
| 40007 | growth | 92 | 8.95E-06 |
| 42180 | cellular ketone metabolic process | 187 | 1.37E-07 |
| 42221 | response to chemical stimulus | 443 | 1.69E-08 |
| 42360 | vitamin E metabolic process | 5 | 4.63E-02 |
| 42362 | fat-soluble vitamin biosynthetic process | 8 | 1.85E-02 |
| 42440 | pigment metabolic process | 37 | 1.94E-04 |
| 42592 | homeostatic process | 59 | 1.12E-02 |
| 42742 | defense response to bacterium | 61 | 1.55E-03 |
| 43412 | macromolecule modification | 312 | 4.24E-02 |
| 43436 | oxoacid metabolic process | 183 | 2.88E-07 |
| 43687 | post-translational protein modification | 260 | 8.87E-03 |
| 44038 | cell wall macromolecule biosynthetic process | 7 | 4.22E-02 |
| 44242 | cellular lipid catabolic process | 23 | 2.42E-03 |
| 44248 | cellular catabolic process | 129 | 2.29E-02 |
| 44255 | cellular lipid metabolic process | 134 | 1.95E-08 |
| 44262 | cellular carbohydrate metabolic process | 112 | 1.23E-02 |
| 44271 | cellular nitrogen compound biosynthetic process | 109 | 4.10E-04 |
| 44281 | small molecule metabolic process | 344 | 1.87E-09 |
| 44282 | small molecule catabolic process | 53 | 1.92E-03 |
| 44283 | small molecule biosynthetic process | 170 | 1.03E-04 |
| 45184 | establishment of protein localization | 103 | 1.38E-05 |
| 45487 | gibberellin catabolic process | 5 | 2.24E-02 |
| 46148 | pigment biosynthetic process | 31 | 6.22E-04 |
| 46394 | carboxylic acid biosynthetic process | 88 | 2.42E-03 |
| 46395 | carboxylic acid catabolic process | 26 | 9.98E-03 |
| 46483 | heterocycle metabolic process | 104 | 5.59E-05 |
| 46486 | glycerolipid metabolic process | 19 | 4.22E-02 |
| 46686 | response to cadmium ion | 93 | 5.10E-06 |
| 46903 | secretion | 17 | 4.41E-02 |
| 46907 | intracellular transport | 109 | 3.44E-07 |
| 46942 | carboxylic acid transport | 25 | 3.96E-02 |
| 48193 | Golgi vesicle transport | 20 | 2.32E-04 |
| 48229 | gametophyte development | 71 | 7.73E-05 |
| 48468 | cell development | 60 | 6.33E-04 |
| 48588 | developmental cell growth | 36 | 2.18E-04 |
| 48589 | developmental growth | 73 | 4.57E-06 |
| 48610 | reproductive cellular process | 45 | 9.18E-04 |
| 48856 | anatomical structure development | 350 | 1.67E-05 |

| | | | |
|---|---|---|---|
| 48868 | pollen tube development | 33 | 4.76E-03 |
| 48869 | cellular developmental process | 130 | 1.59E-05 |
| 48878 | chemical homeostasis | 39 | 5.24E-03 |
| 50801 | ion homeostasis | 29 | 3.70E-02 |
| 50896 | response to stimulus | 794 | 2.97E-11 |
| 51179 | localization | 449 | 9.62E-16 |
| 51186 | cofactor metabolic process | 70 | 3.89E-04 |
| 51188 | cofactor biosynthetic process | 50 | 9.74E-05 |
| 51234 | establishment of localization | 440 | 2.32E-16 |
| 51641 | cellular localization | 134 | 1.09E-07 |
| 51649 | establishment of localization in cell | 126 | 6.83E-08 |
| 51704 | multi-organism process | 207 | 1.69E-08 |
| 51707 | response to other organism | 154 | 8.51E-06 |
| 51716 | cellular response to stimulus | 187 | 1.34E-03 |
| 55046 | microgametogenesis | 10 | 3.03E-02 |
| 55075 | potassium ion homeostasis | 5 | 6.02E-03 |
| 55080 | cation homeostasis | 26 | 2.91E-02 |
| 55086 | nucleobase, nucleoside and nucleotide metabolic process | 54 | 1.49E-02 |
| 55114 | oxidation reduction | 48 | 5.01E-04 |
| 60560 | developmental growth involved in morphogenesis | 63 | 4.73E-05 |
| 60918 | auxin transport | 19 | 2.88E-02 |
| 61024 | membrane organization | 32 | 3.14E-04 |
| 61025 | membrane fusion | 13 | 4.41E-02 |
| 65008 | regulation of biological quality | 174 | 5.10E-08 |
| 70589 | cellular component macromolecule biosynthetic process | 7 | 4.22E-02 |
| 70592 | cell wall polysaccharide biosynthetic process | 7 | 4.22E-02 |
| 70727 | cellular macromolecule localization | 82 | 1.21E-04 |
| 70882 | cellular cell wall organization or biogenesis | 28 | 4.51E-02 |
| 70887 | cellular response to chemical stimulus | 103 | 9.81E-04 |
| 71310 | cellular response to organic substance | 93 | 1.48E-03 |
| 71322 | cellular response to carbohydrate stimulus | 13 | 2.60E-02 |
| 71395 | cellular response to jasmonic acid stimulus | 16 | 2.60E-02 |
| 71495 | cellular response to endogenous stimulus | 73 | 7.05E-03 |
| 90066 | regulation of anatomical structure size | 83 | 4.64E-05 |

#### Supplementary Table S12

| ID | Multimapping | Too Many Loci | Unique |
|---|---|---|---|
| LIB-10X-317-003-1 | 2.93% | 1.61% | 92.67% |
| LIB-10X-317-004-1 | 3.30% | 1.36% | 92.08% |
| LIB-10X-317-005-1 | 2.55% | 13.42% | 78.88% |
| LIB-10X-317-006-1 | 2.83% | 7.18% | 85.96% |
| LIB-10X-317-007-1 | 2.65% | 6.32% | 87.51% |
| LIB-10X-317-008-1 | 2.98% | 11.60% | 81.15% |
| LIB-10X-317-011-1 | 2.88% | 1.12% | 92.50% |
| LIB-10X-317-012-1 | 2.75% | 7.04% | 86.64% |
| LIB-10X-317-013-1 | 2.77% | 11.32% | 81.28% |
| LIB-10X-317-013-2 | 2.82% | 10.65% | 81.57% |
| LIB-10X-317-014-1 | 3.29% | 1.07% | 92.86% |
| LIB-10X-317-014-2 | 3.29% | 1.13% | 92.58% |
| LIB-10X-317-015-1 | 2.79% | 1.62% | 91.85% |
| LIB-10X-317-016-1 | 2.27% | 21.37% | 69.26% |
| LIB-10X-317-017-1 | 2.79% | 0.56% | 93.98% |
| LIB-10X-317-019-1 | 2.72% | 19.54% | 72.35% |
| LIB-10X-317-020-1 | 2.56% | 9.08% | 84.50% |
| LIB-10X-317-020-2 | 2.64% | 8.80% | 84.41% |
| LIB-10X-317-021-1 | 2.75% | 11.15% | 80.39% |
| LIB-10X-317-024-1 | 2.79% | 3.61% | 88.20% |
| LIB-10X-317-028-1 | 3.29% | 3.44% | 89.31% |
| LIB-10X-317-029-1 | 2.61% | 16.51% | 75.58% |
| LIB-10X-317-033-1 | 2.64% | 8.39% | 83.91% |
| LIB-10X-317-034-1 | 2.63% | 18.53% | 72.22% |
| LIB-10X-317-035-1 | 2.59% | 23.00% | 68.72% |
| LIB-10X-317-036-1 | 3.40% | 0.83% | 92.89% |
| LIB-10X-317-036-2 | 3.22% | 0.98% | 93.24% |
| LIB-10X-317-038-1 | 3.07% | 0.71% | 93.71% |
| LIB-10X-317-038-2 | 3.10% | 0.71% | 93.35% |
| LIB-10X-317-040-1 | 2.48% | 16.84% | 75.14% |
| LIB-10X-317-041-1 | 2.67% | 11.94% | 80.70% |
| LIB-10X-317-042-1 | 3.86% | 0.63% | 92.69% |
| LIB-10X-317-044-1 | 2.99% | 1.21% | 92.98% |
| LIB-10X-317-045-1 | 3.02% | 0.49% | 93.51% |
| LIB-10X-317-045-2 | 2.99% | 0.52% | 93.19% |
| LIB-10X-317-046-1 | 2.91% | 0.99% | 93.14% |

| | | | |
|---|---|---|---|
| LIB-10X-317-047-1 | 3.06% | 0.54% | 93.60% |
| LIB-10X-317-047-2 | 3.05% | 0.54% | 93.39% |
| LIB-10X-317-049-1 | 3.44% | 0.78% | 93.31% |
| LIB-10X-317-049-2 | 3.45% | 0.79% | 93.04% |
| LIB-10X-317-050-1 | 2.53% | 19.50% | 73.34% |
| LIB-10X-317-052-1 | 2.47% | 27.04% | 64.34% |
| LIB-10X-317-054-1 | 3.15% | 6.16% | 87.25% |
| LIB-10X-317-055-1 | 2.67% | 8.73% | 83.72% |
| LIB-10X-317-056-1 | 2.68% | 8.79% | 84.12% |
| LIB-10X-317-058-1 | 2.52% | 17.54% | 74.28% |
| LIB-10X-317-059-1 | 2.51% | 16.40% | 74.72% |
| LIB-10X-317-060-1 | 3.21% | 1.11% | 92.87% |
| LIB-10X-317-061-1 | 2.94% | 0.56% | 93.81% |
| LIB-10X-317-065-1 | 2.79% | 0.97% | 93.43% |
| LIB-10X-317-068-1 | 2.73% | 16.15% | 74.81% |
| LIB-10X-317-069-1 | 2.88% | 0.94% | 92.21% |
| LIB-10X-317-076-1 | 2.85% | 7.84% | 84.68% |
| LIB-10X-317-078-1 | 3.29% | 3.57% | 89.15% |
| LIB-10X-317-081-1 | 2.92% | 2.83% | 90.50% |
| LIB-10X-317-086-1 | 2.75% | 10.52% | 82.37% |
| LIB-10X-317-089-1 | 3.10% | 2.25% | 90.57% |
| LIB-10X-317-091-1 | 3.08% | 9.09% | 83.83% |
| LIB-10X-317-092-1 | 2.89% | 4.55% | 88.81% |
| LIB-10X-317-093-1 | 2.81% | 12.20% | 79.42% |
| LIB-10X-317-097-1 | 1.86% | 34.16% | 54.41% |
| LIB-10X-317-099-1 | 1.67% | 39.24% | 48.79% |
| LIB-10X-317-100-1 | 1.68% | 51.67% | 35.56% |
| LIB-10X-317-101-1 | 1.99% | 35.71% | 51.83% |
| LIB-10X-317-102-1 | 2.99% | 0.91% | 93.11% |
| LIB-10X-317-104-1 | 2.69% | 16.02% | 75.67% |
| LIB-10X-317-105-1 | 2.91% | 1.66% | 92.27% |
| LIB-10X-317-106-1 | 2.84% | 0.60% | 93.82% |
| LIB-10X-317-107-1 | 3.38% | 0.81% | 92.93% |
| LIB-10X-317-108-1 | 3.06% | 1.35% | 92.36% |
| LIB-10X-317-109-1 | 2.65% | 10.98% | 80.92% |
| LIB-10X-317-112-1 | 2.69% | 13.61% | 78.08% |
| LIB-10X-317-113-1 | 3.36% | 0.57% | 92.13% |
| LIB-10X-317-114-1 | 1.81% | 45.14% | 41.19% |
| LIB-10X-317-115-1 | 2.69% | 19.43% | 72.65% |
| LIB-10X-317-117-1 | 3.06% | 0.55% | 93.43% |

| | | | |
|---|---|---|---|
| LIB-10X-317-120-1 | 3.02% | 0.82% | 93.33% |
| LIB-10X-317-121-1 | 3.28% | 0.58% | 93.39% |
| LIB-10X-317-122-1 | 2.76% | 2.82% | 91.54% |
| LIB-10X-317-124-1 | 3.12% | 0.49% | 93.16% |
| LIB-10X-317-127-1 | 3.70% | 1.02% | 92.79% |
| LIB-10X-317-127-2 | 3.72% | 1.01% | 92.45% |
| LIB-10X-317-129-1 | 2.74% | 6.46% | 87.30% |
| LIB-10X-317-134-1 | 1.47% | 42.78% | 46.96% |
| LIB-10X-317-134-2 | 1.56% | 41.45% | 47.80% |
| LIB-10X-317-135-1 | 3.05% | 0.74% | 92.20% |
| LIB-10X-317-135-2 | 3.11% | 0.70% | 91.71% |
| LIB-10X-317-136-1 | 2.87% | 4.91% | 89.45% |
| LIB-10X-317-136-2 | 2.91% | 4.99% | 89.06% |
| LIB-10X-317-138-1 | 1.77% | 36.66% | 51.84% |
| LIB-10X-317-139-1 | 2.09% | 32.08% | 58.56% |
| LIB-10X-317-139-2 | 2.14% | 31.37% | 58.73% |
| LIB-10X-317-140-1 | 3.01% | 5.40% | 88.05% |
| LIB-10X-317-140-2 | 3.06% | 5.12% | 88.04% |
| LIB-10X-317-141-1 | 1.88% | 36.45% | 52.31% |
| LIB-10X-317-145-1 | 1.86% | 37.11% | 53.00% |
| LIB-10X-317-145-2 | 1.83% | 38.06% | 51.64% |
| LIB-10X-317-146-1 | 3.44% | 2.08% | 91.68% |
| LIB-10X-317-146-2 | 3.38% | 2.30% | 91.32% |
| LIB-10X-317-147-1 | 2.80% | 1.03% | 92.58% |
| LIB-10X-317-149-1 | 2.89% | 1.55% | 91.86% |
| LIB-10X-317-152-1 | 2.99% | 1.24% | 92.19% |
| LIB-10X-317-153-1 | 2.99% | 0.73% | 92.74% |
| LIB-10X-317-155-1 | 3.29% | 1.25% | 92.63% |
| LIB-10X-317-155-2 | 3.31% | 1.23% | 92.37% |
| LIB-10X-317-156-1 | 3.04% | 0.87% | 93.62% |
| LIB-10X-317-159-1 | 2.54% | 17.20% | 74.47% |
| LIB-10X-317-159-2 | 2.54% | 17.32% | 74.25% |
| LIB-10X-317-161-1 | 2.72% | 7.06% | 85.11% |
| LIB-10X-317-161-2 | 2.76% | 7.06% | 84.53% |
| LIB-10X-317-163-1 | 2.86% | 0.98% | 93.64% |
| LIB-10X-317-164-1 | 3.35% | 2.86% | 91.16% |
| LIB-10X-317-164-2 | 3.33% | 3.09% | 90.70% |
| LIB-10X-317-165-1 | 2.92% | 6.09% | 87.39% |
| LIB-10X-317-165-2 | 2.92% | 6.80% | 86.36% |
| LIB-10X-317-166-1 | 3.04% | 0.70% | 93.86% |

| | | | |
|---|---|---|---|
| LIB-10X-317-168-1 | 2.75% | 9.21% | 82.70% |
| LIB-10X-317-171-1 | 2.92% | 0.44% | 94.07% |
| LIB-10X-317-175-1 | 2.01% | 31.61% | 56.89% |
| LIB-10X-317-176-1 | 3.05% | 3.43% | 90.39% |
| LIB-10X-317-178-1 | 3.31% | 0.74% | 93.24% |
| LIB-10X-317-178-2 | 3.28% | 0.80% | 92.83% |
| LIB-10X-317-181-1 | 2.94% | 1.25% | 92.52% |
| LIB-10X-317-182-1 | 3.15% | 1.05% | 93.31% |
| LIB-10X-317-183-1 | 2.91% | 1.34% | 93.31% |
| LIB-10X-317-184-1 | 3.05% | 1.82% | 91.79% |
| LIB-10X-317-186-1 | 3.34% | 0.70% | 92.74% |
| LIB-10X-317-190-1 | 2.91% | 1.75% | 92.92% |
| LIB-10X-317-191-1 | 3.41% | 0.63% | 93.38% |
| LIB-10X-317-191-2 | 3.45% | 0.66% | 93.00% |
| LIB-10X-317-192-1 | 2.30% | 19.98% | 71.46% |
| LIB-10X-317-195-1 | 1.85% | 33.92% | 56.66% |
| LIB-10X-317-195-2 | 1.89% | 33.82% | 56.23% |
| LIB-10X-317-197-1 | 2.54% | 12.70% | 79.33% |
| LIB-10X-317-198-1 | 2.44% | 19.68% | 72.16% |
| LIB-10X-317-198-2 | 2.50% | 19.26% | 72.19% |
| LIB-10X-317-199-1 | 2.76% | 9.96% | 82.82% |
| LIB-10X-317-201-1 | 2.80% | 5.13% | 88.66% |
| LIB-10X-317-203-1 | 3.22% | 0.89% | 92.55% |
| LIB-10X-317-203-2 | 3.29% | 0.90% | 92.02% |
| LIB-10X-317-204-1 | 2.68% | 13.71% | 78.65% |
| LIB-11X-317-002-1 | 2.03% | 24.80% | 64.91% |
| LIB-11X-317-003-1 | 2.71% | 6.96% | 86.56% |
| LIB-11X-317-003-2 | 2.69% | 7.09% | 86.28% |
| LIB-11X-317-004-1 | 3.16% | 0.57% | 93.71% |
| LIB-11X-317-004-2 | 3.20% | 0.58% | 93.34% |
| LIB-11X-317-008-1 | 3.16% | 3.76% | 90.58% |
| LIB-11X-317-008-2 | 3.21% | 3.92% | 90.08% |
| LIB-11X-317-009-1 | 3.18% | 0.64% | 93.10% |
| LIB-11X-317-009-2 | 3.24% | 0.63% | 92.74% |
| LIB-11X-317-012-1 | 2.69% | 8.02% | 85.20% |
| LIB-11X-317-012-2 | 2.69% | 8.79% | 83.83% |
| LIB-11X-317-018-1 | 2.31% | 17.40% | 74.18% |
| LIB-11X-317-018-2 | 2.31% | 17.83% | 73.56% |
| LIB-11X-317-022-1 | 2.47% | 21.40% | 68.65% |
| LIB-11X-317-022-2 | 2.47% | 21.56% | 68.33% |

| | | | |
|---|---|---|---|
| LIB-11X-317-024-1 | 2.44% | 17.43% | 73.76% |
| LIB-11X-317-024-2 | 2.45% | 17.64% | 73.39% |
| LIB-11X-317-029-1 | 3.17% | 9.37% | 83.75% |
| LIB-11X-317-029-2 | 3.07% | 11.00% | 81.62% |
| LIB-11X-317-030-1 | 2.76% | 3.87% | 88.86% |
| LIB-11X-317-030-2 | 2.74% | 3.92% | 88.66% |
| LIB-11X-317-034-1 | 2.60% | 12.10% | 78.56% |
| LIB-11X-317-034-2 | 2.59% | 12.15% | 77.98% |
| LIB-11X-317-039-1 | 3.28% | 1.09% | 92.38% |
| LIB-11X-317-039-2 | 3.21% | 1.11% | 91.87% |
| LIB-11X-317-042-1 | 2.96% | 1.22% | 92.85% |
| LIB-11X-317-042-2 | 2.95% | 1.27% | 92.41% |
| LIB-11X-317-043-1 | 2.56% | 19.07% | 72.58% |
| LIB-11X-317-043-2 | 2.55% | 19.69% | 71.47% |
| LIB-11X-317-046-1 | 3.03% | 2.75% | 91.63% |
| LIB-11X-317-049-1 | 2.81% | 0.51% | 93.50% |
| LIB-11X-317-051-1 | 3.11% | 5.73% | 86.89% |
| LIB-11X-317-051-2 | 2.95% | 6.52% | 86.01% |
| LIB-11X-317-055-1 | 1.57% | 46.01% | 40.48% |
| LIB-11X-317-058-1 | 3.16% | 0.59% | 92.75% |
| LIB-11X-317-058-2 | 3.18% | 0.59% | 92.29% |
| LIB-11X-317-060-1 | 2.38% | 22.79% | 68.14% |
| LIB-11X-317-060-2 | 2.42% | 22.27% | 68.33% |
| LIB-11X-317-063-1 | 1.84% | 40.15% | 49.93% |
| LIB-11X-317-063-2 | 1.90% | 40.36% | 49.19% |
| LIB-11X-317-064-1 | 2.74% | 7.83% | 84.82% |
| LIB-11X-317-068-1 | 2.06% | 37.19% | 50.32% |
| LIB-11X-317-069-1 | 3.25% | 0.61% | 93.54% |
| LIB-11X-317-069-2 | 3.27% | 0.63% | 93.25% |
| LIB-11X-317-075-1 | 3.17% | 0.93% | 93.69% |
| LIB-11X-317-075-2 | 3.18% | 0.90% | 93.55% |
| LIB-11X-317-076-1 | 3.05% | 2.78% | 90.90% |
| LIB-11X-317-076-2 | 3.11% | 2.69% | 90.72% |
| LIB-11X-317-080-1 | 3.62% | 0.54% | 93.79% |
| LIB-11X-317-083-1 | 2.77% | 18.75% | 73.25% |
| LIB-11X-317-084-1 | 2.85% | 0.57% | 94.13% |
| LIB-11X-317-086-1 | 3.20% | 0.61% | 94.24% |
| LIB-11X-317-089-1 | 2.95% | 0.74% | 94.04% |
| LIB-11X-317-093-1 | 3.41% | 0.91% | 92.96% |
| LIB-11X-317-103-1 | 1.86% | 41.64% | 48.34% |

| | | | |
|---|---|---|---|
| LIB-11X-317-106-1 | 2.99% | 0.66% | 93.24% |
| LIB-11X-317-108-1 | 3.44% | 0.56% | 93.65% |
| LIB-11X-317-110-1 | 2.92% | 0.80% | 93.87% |
| LIB-11X-317-116-1 | 2.87% | 2.73% | 91.31% |
| LIB-11X-317-117-1 | 3.24% | 0.69% | 93.55% |
| LIB-11X-317-118-1 | 2.91% | 0.92% | 93.22% |
| LIB-11X-317-123-1 | 2.99% | 3.46% | 90.71% |
| LIB-11X-317-128-1 | 3.16% | 0.62% | 92.53% |
| LIB-11X-317-130-1 | 3.27% | 0.55% | 93.77% |
| LIB-11X-317-135-1 | 2.81% | 17.51% | 74.51% |
| LIB-11X-317-190-1 | 2.95% | 11.92% | 80.44% |
| LIB-11X-317-193-1 | 2.68% | 14.05% | 78.49% |
| LIB-11X-317-194-1 | 2.88% | 1.86% | 91.94% |
| LIB-11X-317-197-1 | 3.00% | 1.46% | 93.18% |
| LIB-11X-317-198-1 | 2.95% | 0.80% | 93.10% |
| LIB-11X-317-199-1 | 2.45% | 16.44% | 74.44% |
| LIB-11X-317-200-1 | 2.91% | 0.97% | 92.72% |
| LIB-11X-317-203-1 | 2.72% | 9.07% | 83.52% |
| LIB-11X-317-204-1 | 2.73% | 1.05% | 92.61% |
| LIB-11X-317-205-1 | 2.51% | 15.09% | 75.97% |
| LIB-11X-317-208-1 | 2.65% | 7.98% | 84.21% |
| LIB-11X-317-212-1 | 2.26% | 17.74% | 72.69% |
| LIB-11X-317-213-1 | 3.08% | 4.27% | 90.21% |
| LIB-11X-317-217-1 | 1.76% | 34.25% | 54.12% |
| LIB-11X-317-231-1 | 2.65% | 8.98% | 83.30% |
| LIB-11X-317-232-1 | 2.74% | 2.08% | 91.69% |
| LIB-11X-317-234-1 | 2.81% | 0.61% | 93.83% |
| LIB-11X-317-239-1 | 2.87% | 2.58% | 90.93% |
| LIB-11X-317-240-1 | 2.81% | 4.77% | 88.02% |
| LIB-94006-1 | 3.34% | 8.37% | 85.73% |

**Supplementary Figure S1**

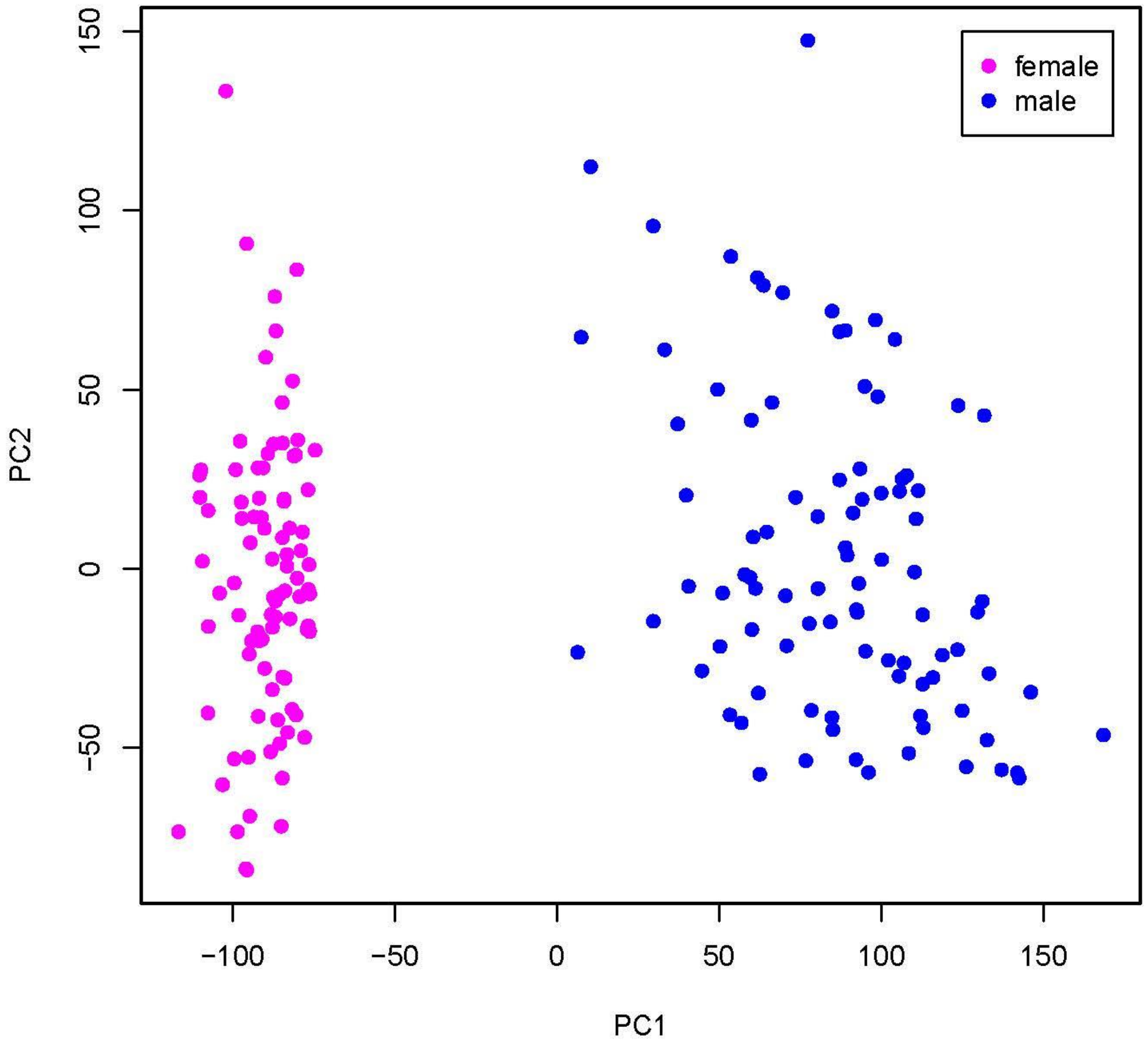

**Supplementary Figure S2**

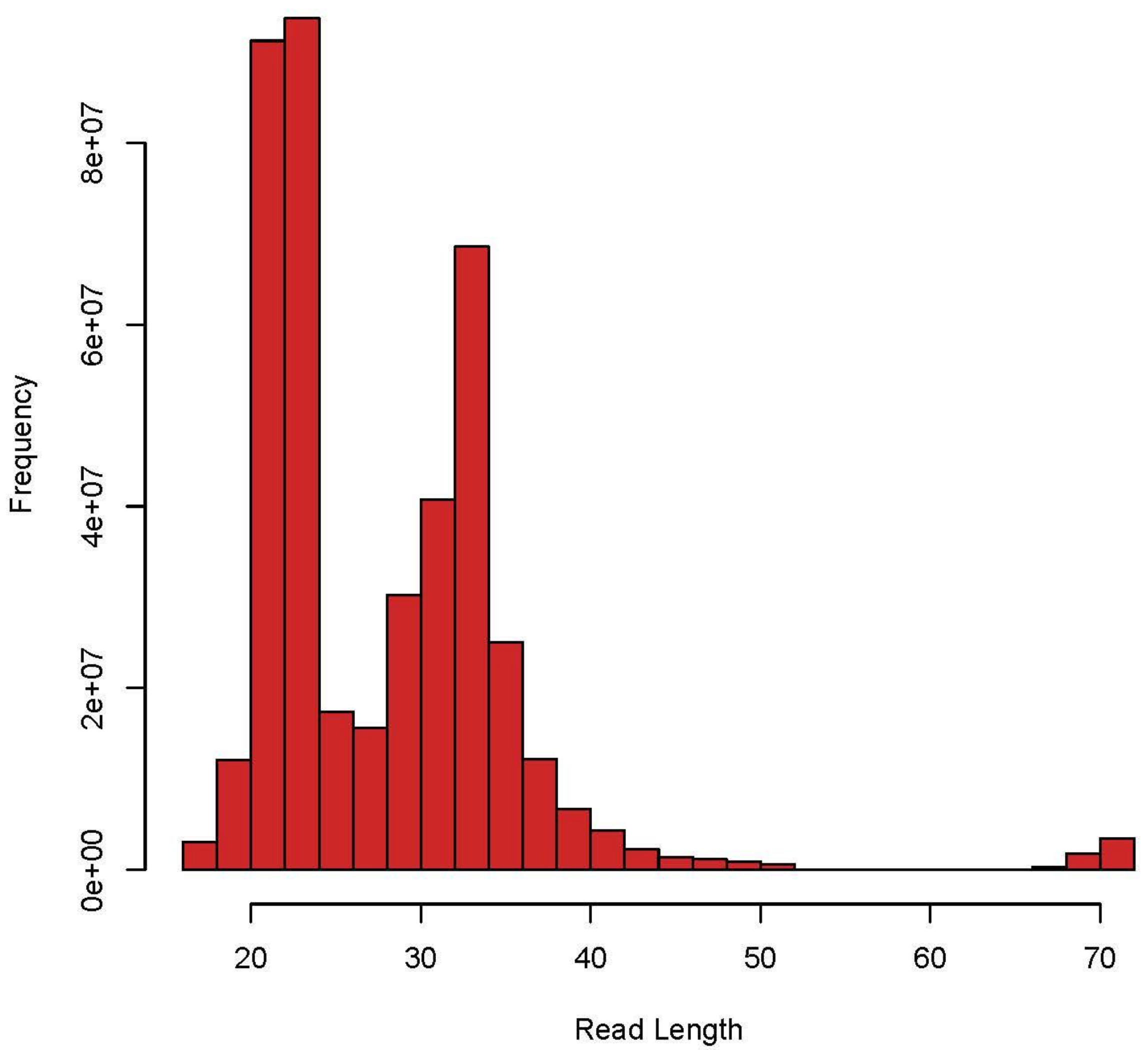

#### Supplementary Figure S4A

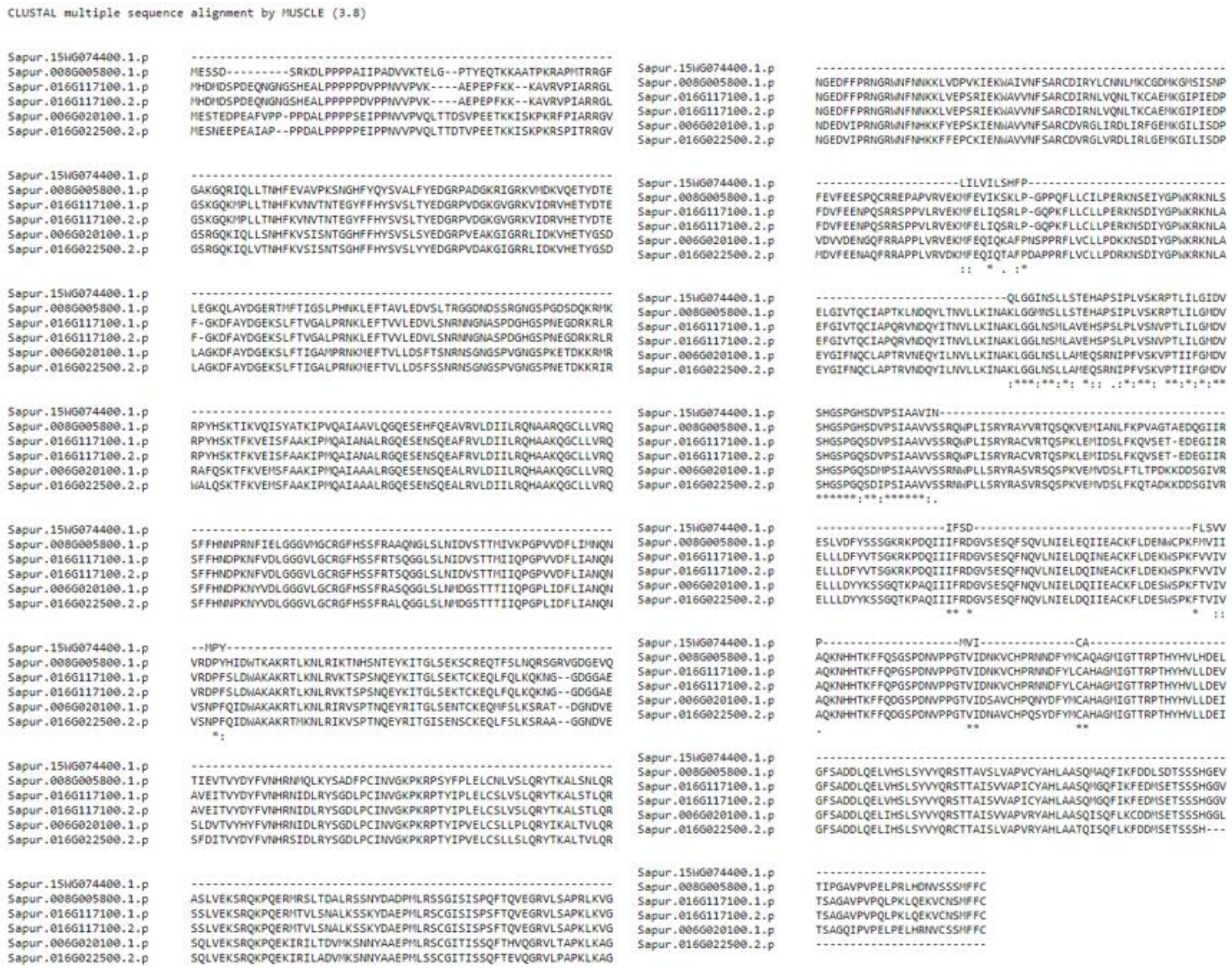


#### Supplementary Figure S4B

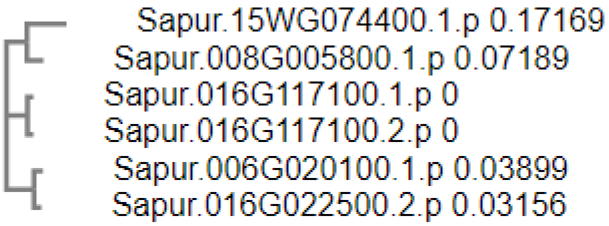

### Supplementary Figure S5A

```
Cluster_187938 Original Location: Chr15Z:5200310-5200523 Displayed Location: Chr15Z:5200405-5200526 Strand: +

GGAGGCACUGAUGAUGUUGUUGACAGAAGAUAGAGAGCACAGAUGAUGGUAUGCAAUGGGCUCUGCAUCCCACUCCUUUGUGCUCUCUAAGCUUCUGUCAUCACUUUCAGCCCCCGACCCCC
((.(((.....(((...((.(((((((((.(((((((((((((.(((((.(((((........))))).))).)).)))))))))))))..))))))))).))...)))))).))..)))))
.................................................................................GCUCUCUAAGCUUCUGUCAUC.................... miRNA l=21 a=1487
....................UGACAGAAGAUAGAGAGCAC.................................................................................. miRNA-star l=20 a=4

....................................................................................CUCUAAGCUUCUGUCAUCc................... l=19 a=1
...................................................................................UCUCUAAGCUUCUGUCAUC.................... l=19 a=2
...................................................................................UCUCUAAGCUUCUGUCAUCACU................. l=22 a=5
..................................................................................CUCUCUAAGCUUCUGUCAUC.................... l=20 a=2
..................................................................................CUCUCUAAGCUUCUGUCAUCA................... l=21 a=1
..................................................................................CUCUCUAAGCUUCUGUCAUCACUUUCAGCCCCCGACu... l=37 a=2
..................................................................................CUCUCUAuGCUUCUGUCAUC.................... l=20 a=1
.................................................................................GCUCUCUAAGCUUCUGUC....................... l=18 a=1
.................................................................................GCUCUCUAAGCUUCUGUCA...................... l=19 a=1
.................................................................................GCUCUCUAAGCUUCUGUCAU..................... l=20 a=4
.................................................................................GCUCUCUAAGCUUCUGUCAUCA................... l=22 a=19
.................................................................................GCUCUCUAAGCUUCUGUCAUCACUUUCAGCCCCCGACu... l=38 a=1
.................................................................................GCUCUCUAAGCUUCUGUCAUCc................... l=22 a=3
.................................................................................GCUCUCUAAGCUUCUGUCAUCu................... l=22 a=98
.................................................................................GCUCUCUAAGCUUCUGUCAUCuC.................. l=23 a=1
.................................................................................GCUCUCUAAGCUUCUGUCAUa.................... l=21 a=4
.................................................................................GCUCUCUAAGCUUCUGUCAUu.................... l=21 a=36
.................................................................................GCUCUCUAAGCUUCUGUaAUC.................... l=21 a=4
.................................................................................GCUCUCUAAGCUUCUGaCAUC.................... l=21 a=1
.................................................................................GCUCUCUAAGCUUCUuUCAUC.................... l=21 a=1
.................................................................................GCUCUCUAAGCUUCaGUCAUC.................... l=21 a=3
.................................................................................GCUCUCUAAGCUUCcGUCAUC.................... l=21 a=1
.................................................................................GCUCUCUAAGCUUCgGUCAUC.................... l=21 a=1
.................................................................................GCUCUCUAAGCUUaUGUCAUC.................... l=21 a=1
.................................................................................GCUCUCUAAGCUUaUGUCAUCA................... l=22 a=1
.................................................................................GCUCUCUAAGCcUCUGUCAUC.................... l=21 a=1
.................................................................................GCUCUCUAAGaUUCUGUCAUC.................... l=21 a=4
.................................................................................GCUCUCUAuGCUUCUGUCAUC.................... l=21 a=77
.................................................................................GCUCUCUAuGCUUCUGUCAUCA................... l=22 a=3
.................................................................................GCUCUCaAAGCUUCUGUCAUC.................... l=21 a=1
.................................................................................GCUCUaUAAGCUUCUGUCAUC.................... l=21 a=1
.................................................................................GCUCUuUAAGCUUCUGUCAUC.................... l=21 a=1
.................................................................................GCUCaCUAAGCUUCUGUCAUC.................... l=21 a=2
.................................................................................GCUaUCUAAGCUUCUGUCAUC.................... l=21 a=1
.................................................................................GgUCUCUAAGCUUCUGUCAUC.................... l=21 a=2
.................................................................................cCUCUCUAAGCUUCUGUCAUC.................... l=21 a=1
.................................................................................nCUCUCUAAGCUUCUGUCAUC.................... l=21 a=1
.................................................................................uCUCUCUAAGCUUCUGUCAUC.................... l=21 a=5
................................................................................UGCUCUCUAAGCUUCUGUCAUC.................... l=22 a=1
........................................................................CUCCUUUGUGCUCUCUAAGCUU............................ l=22 a=5
.....................................................................CCACUCCUUUGUGCUCUCUAAGCU............................. l=24 a=1
.....................................................................CCACUCCUUUGUGCUCUCUAAGCUU............................ l=25 a=2
.................................................................CAUCCCACUCCUUUGUGCUCUCUAAGCUUCU.......................... l=31 a=1
........................................AGAUGAUGGUAUGCAAUGGG.............................................................. l=20 a=1
........................................AGAUGAUGGUAUGCAAUGGGC............................................................. l=21 a=2
........................................AGAUGAUGGUAUGCAAUGGGCU............................................................ l=22 a=1
........................................AGAUGAUGGUAUGCAAUGGGCUCUGCAUCC.................................................... l=30 a=1
........................................AGAUGAUGGUAUGCAAUGGGCUCUGCAUCCC................................................... l=31 a=3
........................................AGAUGAUGGUAUGCAAUGGGCUCUGCAUCCCA.................................................. l=32 a=1
........................................AGAUGAUGGUAUGCAAUGGGCUCUGCAUCCCACUCCUUUGU......................................... l=41 a=2
...................................AGCACAGAUGAUGGUAUGCAAUGGGC............................................................. l=26 a=1
............................GAUAGAGAGCACAGAUGAUGG......................................................................... l=21 a=2
.....................GACAGAAGAUAGAGAGCAC.................................................................................. l=19 a=1
....................UGACAGAAGAUAGAGAGCACAc................................................................................ l=22 a=1
...................UUGACAGAAGAUAGAGAGC.................................................................................... l=19 a=22
...................UUGACAGAAGAUAGAGAGCA................................................................................... l=20 a=8
...................UUGACAGAAGAUAGAGAGCAC.................................................................................. l=21 a=611
...................UUGACAGAAGAUAGAGAGCACA................................................................................. l=22 a=2
...................UUGACAGAAGAUAGAGAGCACAc................................................................................ l=23 a=1
...................UUGACAGAAGAUAGAGAGCACAu................................................................................ l=23 a=26
...................UUGACAGAAGAUAGAGAGCACc................................................................................. l=22 a=6
...................UUGACAGAAGAUAGAGAGCACu................................................................................. l=22 a=247
...................UUGACAGAAGAUAGAGAGCAaA................................................................................. l=22 a=1
...................UUGACAGAAGAUAGAGAGCAu.................................................................................. l=21 a=1
...................UUGACAGAAGAUAGAGAGCAuA................................................................................. l=22 a=7
...................UUGACAGAAGAUAGAGAGCu................................................................................... l=20 a=1
...................UUGACAGAAGAUAGAGgGCAC.................................................................................. l=21 a=1
...................UUGACAGAAcAUAGAGAGCACA................................................................................. l=22 a=1
...................UUGAgAGAAGAUAGAGAGCACA................................................................................. l=22 a=1
...................UUGgCAGAAGAUAGAGAGCAC.................................................................................. l=21 a=2
...................UUuACAGAAGAUAGAGAGCACA................................................................................. l=22 a=1
...................UaGACAGAAGAUAGAGAGCAC.................................................................................. l=21 a=1
...................UaGACAGAAGAUAGAGAGCACA................................................................................. l=22 a=3
...................UcGACAGAAGAUAGAGAGCAC.................................................................................. l=21 a=1
...................aUGACAGAAGAUAGAGAGCAC.................................................................................. l=21 a=1
................UUGUUGACAGAAGAUAGAGAGCA................................................................................... l=23 a=1
........UGAUGAUGUUGUUGACAGAAGAUAGAGAGCACA................................................................................. l=33 a=1
.....CACUGAUGAUGUUGUUGACAGAAGAUAGAGAGCAC.................................................................................. l=35 a=2
..AGGCACUGAUGAUGUUGUUGACAGAA.............................................................................................. l=26 a=2
..AGGCACUGAUGAUGUUGUUGACAGAAGAUAGAGAGCA................................................................................... l=37 a=1
<<<<<<<<<<<<<<<<<<<<<<<<<<<<<<<<<<<<<<<<<<<<<<<<<<<<<<<<<<<<<<<<<<<<<<<<<<<<<<<<cCGAGAGAUUCGAAGACAGUA<<<<<<<<<<<<<<<<<<<<< l=21 a=1
<<<<<<<<<<<<<<<<<<<<<<<<<<<<<<<<<<<<<<<<<<<<<<<<<<<<<<<<<<<<<<<<<<<<<<<<<<<<<<<CuCGAGAGAUUCGAAGACAGUA<<<<<<<<<<<<<<<<<<<<< l=22 a=1
<<<<<<<<<<<<<<<<<<<<<<<<<<<<<<<<<<<<<<<<<<<<<<<<<<<<<<<<<<<CGAGACGUAGGGUGAGGAAA<<<<<<<<<<<<<<<<<<<<<<<<<<<<<<<<<<<<<<<<<<< l=20 a=1
<<<<<<<<<<<<<<<<<<<<<<<<<<<<<<<<<<<<<<<<<<<<<<<<<<<<<<<<<<CCGAGACGUAGGGUGAGGAAA<<<<<<<<<<<<<<<<<<<<<<<<<<<<<<<<<<<<<<<<<<< l=21 a=1
<<<<<<<<<<<<<<<<<<<<<<<<<<<<<<<<<<<<<<<<<<<<<<<<<<<<<<<<<CCCGAGACGUAGGGUGAGGAAA<<<<<<<<<<<<<<<<<<<<<<<<<<<<<<<<<<<<<<<<<<< l=22 a=1
```

#### Supplementary Figure S5B

```
 Score = 212 bits (234),  Expect = 4e-54
 Identities = 121/122 (99%), Gaps = 1/122 (1%)
 Strand=Plus/Minus

Query  1        GGAGGCACTGATGATGTTGTTGACAGAAGATAGAGAGCACAGATGATGGTATGCAATGGG  60
                ||||||||||||||||||||||||||||||||||||||||||||||||||||||||||||
Sbjct  4026937  GGAGGCACTGATGATGTTGTTGACAGAAGATAGAGAGCACAGATGATGGTATGCAATGGG  4026878

Query  61       CTCTGCATCCCACTCCTTTGTGCTCTCTAAGCTTCTGTCATCACTTTCAGCCCCCGACCC  120
                |||||||||||||||||||||||||||||||||||||||||||||||||| ||||||||||
Sbjct  4026877  CTCTGCATCCCACTCCTTTGTGCTCTCTAAGCTTCTGTCATCACTTTCAG-CCCCGACCC  4026819

Query  121      CC  122
                ||
Sbjct  4026818  CC  4026817
```

**Supplementary Figure S6**

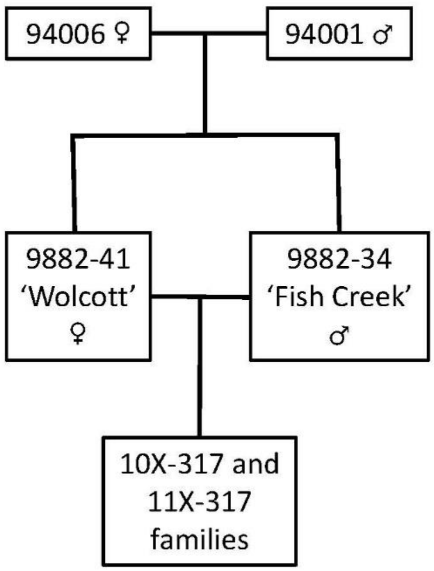

#### Supplementary Figure S7

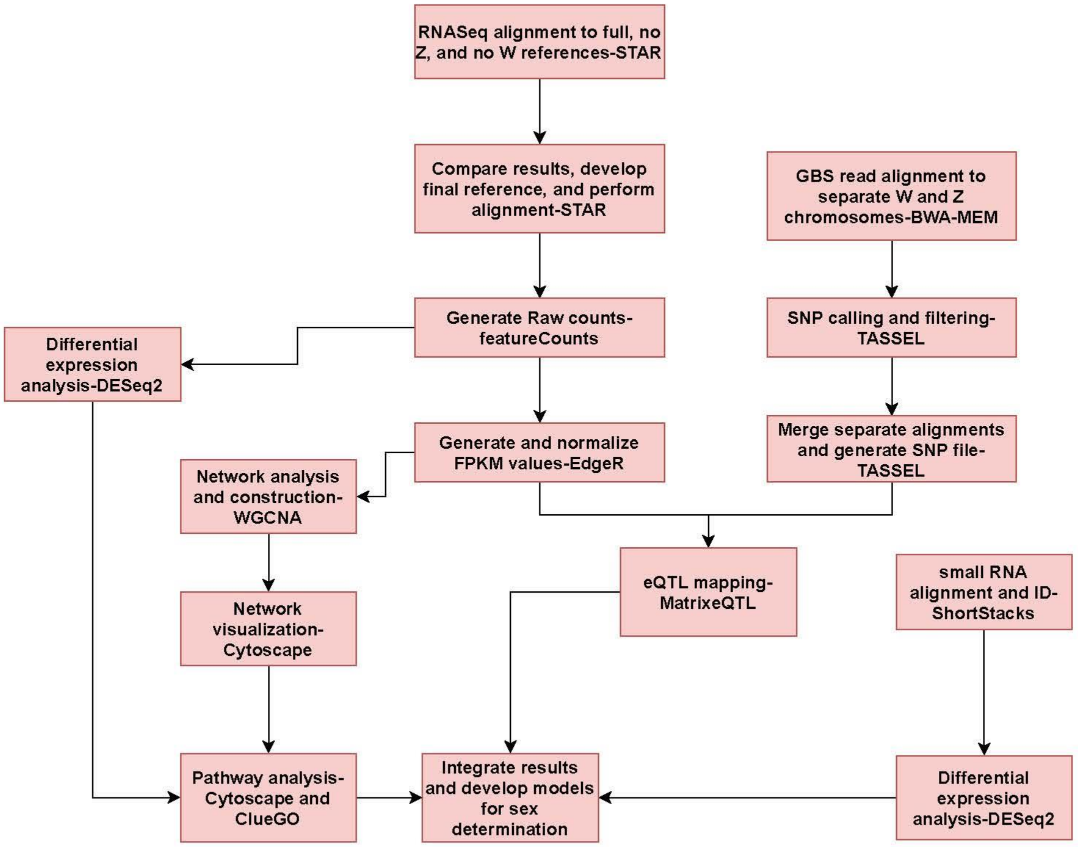


#### Supplementary Figure S8

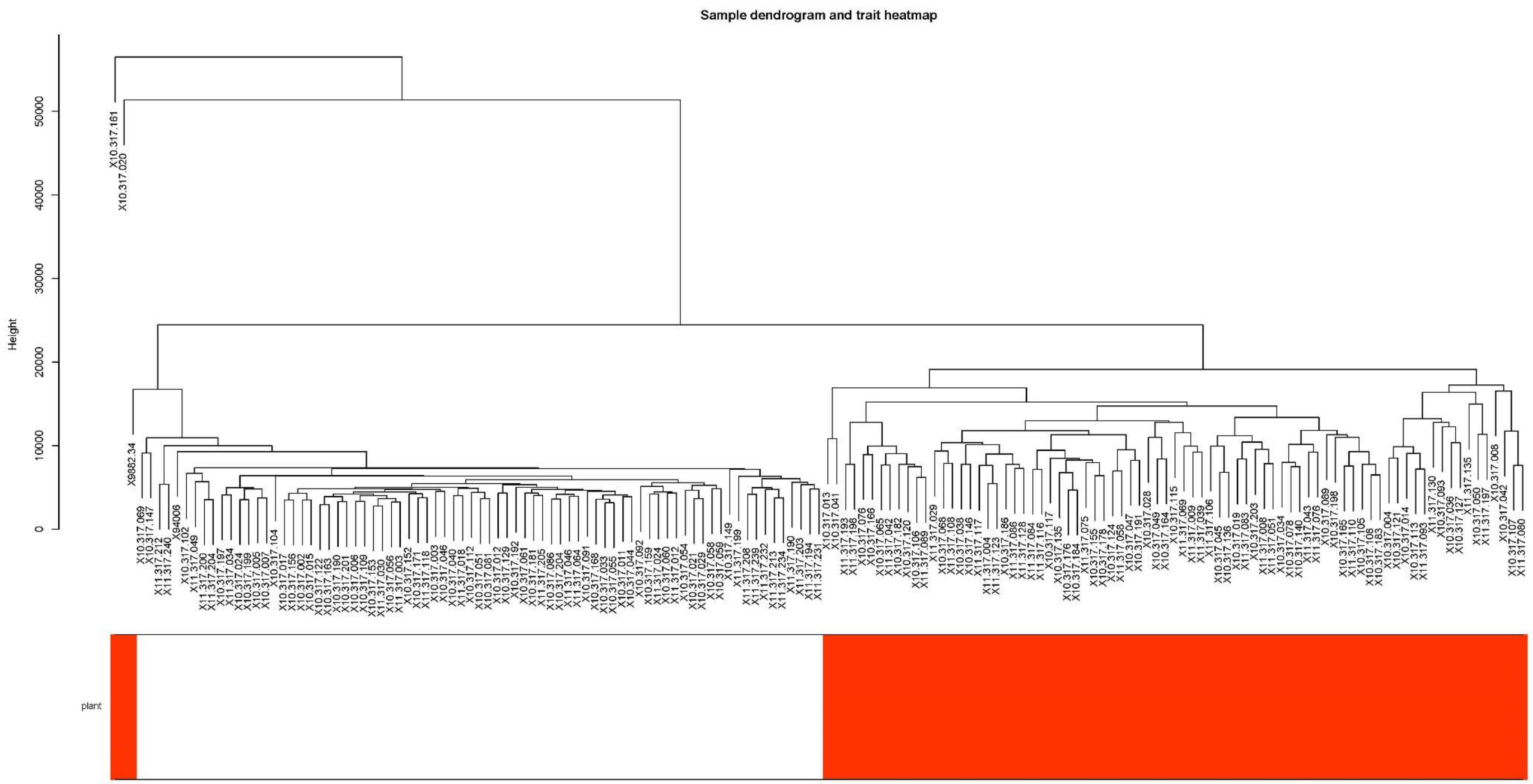

Sample dendrogram and trait heatmap
Height
0
10000
20000
30000
40000
50000
plant
