## Supplementary figures and images for "Integrative genomics reveals paths to sex dimorphism in *Salix purpurea* L."

### Supplementary Fig. S3

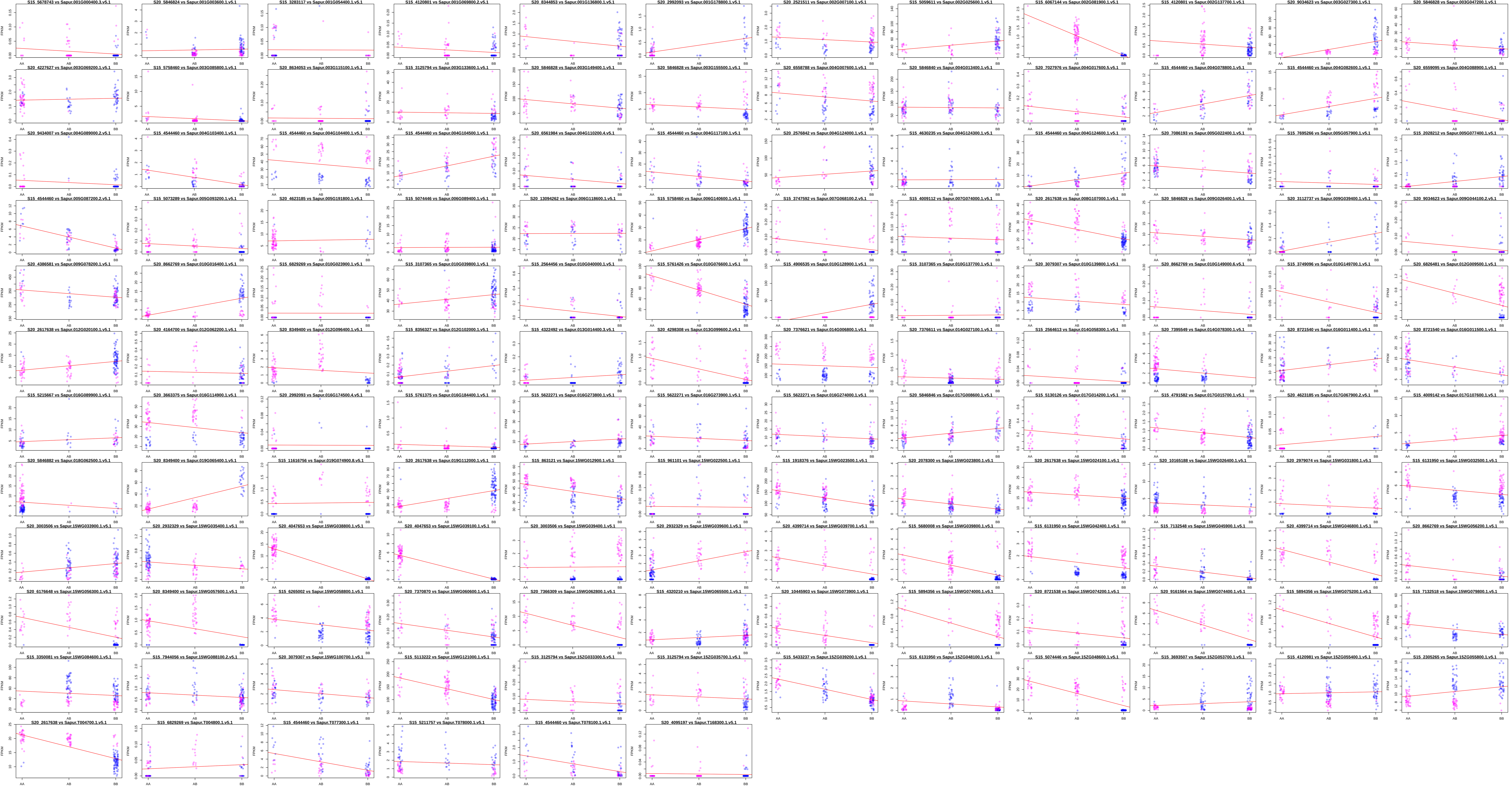
