## Supplementary Methods for "Integrative genomics reveals paths to sex dimorphism in *Salix purpurea* L."

### Materials and Methods

#### *Phenological stage and collection of catkin inflorescences*

After generative bud burst, floral phenology was visually scored on a daily basis, enabling the capture of differentiation prior to receptivity and anthesis in males and females, respectively. Applying the Biologishe Bundesanstalt, Bundessortenamt and Chemical Industry codec, Saska and Kuzovkina (2010) denote this phenological stage as between inflorescence maximum expansion (59) and the onset of flowering (61).

#### *Total RNA-Seq*

Emerging catkins were collected between 10 AM and 2 PM from second-year post-coppice field grown stems of $F_2$ progeny male and female individuals and their parents. At least three catkins each of 90 $F_2$ males and 90 $F_2$ females were flash-frozen in liquid nitrogen immediately after excision from donor plants and stored in 50 mL conical tubes at −80°C, prior to RNA isolation.

For each genotype, a single representative catkin, which contains hundreds of individual flower across a range of developmental stages, was removed from −80°C storage, then ground to a fine powder for RNA isolation using the Spectrum™ Total Plant RNA Kit with DNase I digestion (Sigma, St. Louis, MO). The manufacturer's protocol was followed, with the exception that prior to the tissue lysis step, the 2-ME/lysate mixture was incubated at 65°C for 5 min. Cold ethanol precipitations were performed by the addition of 10 μL acetic acid and 280 μL 100% cold ethanol to 100 μL eluate and placed in −80°C for 3 h. Samples were centrifuged at 17,000 x g for 30 min at 4 °C, washed with 80% ethanol, then centrifuged at 17,000×*g* for 20 min at 4°C. After centrifugation, the supernatant was discarded, and the pellet resuspended in ribonuclease-free 10 mM Tris-HCl. Quantification of RNA sample quality and concentration was performed using the Experion 'StdSens' kit (Bio-Rad Laboratories, Inc., Hercules, CA). Stranded RNA-Seq libraries were created and quantified by qPCR. Paired-end (2×151) sequencing was performed on an Illumina Hi-Seq 2500 system at the Department of Energy Joint Genome Institute (Berkeley, CA).

Reads were aligned to the *S. purpurea* 94006 v5.1 reference genome using STAR, which includes fully assembled and annotated 15Z and 15W sex chromosomes. Chromosomes 15Z and 15W have very low sequence differentiation for most genes, particularly in the psuedoautosomal regions but with many homologous genes across the SDR as well, which causes inaccurate read alignments when both sequences are included in the reference. To circumvent this problem, we performed alignments for references with and without chr15Z. If the read alignments were significantly different based on a Student's T test, the chr15Z allele was included in the reference genome. The final reference therefore included all of chr15W plus 187 genes with flanking sequence from chr15Z. Alignment of RNA-Seq reads to the final reference genome using STAR yielded an average unique mapping rate of 83.45% across samples (table S12). RNA-Seq data from 94006, the individual used to create the reference genome, had a unique mapping rate of 85.73%, and provided a standard for assessing the mapping quality of the F2 individuals. Samples with less than 70% unique read mapping were removed from further analysis, resulting in a final total of 159 individuals, including 77 females and 82 males with an average unique read mapping rate of 87.65%. Raw counts were assigned to mapped reads using featureCounts and utilized to conduct differential expression analysis using the R package DESeq2 in R version 3.5.1. Log2-fold change of male to female expression and FDR adjusted p-values were calculated for each transcript. FPKM values were calculated in the edgeR package for use in subsequent network and eQTL analyses. The resulting data were filtered to remove any genes present in fewer than twenty samples, resulting in 33,880 gene models (including alternative transcripts) being included for final analysis.

*Small RNA-Seq*

For smRNA-Seq, a total of 22 male and 21 female $F_2$ progeny individuals, the male (94001) and female (94006) grandparents of the $F_2$ were collected using the same protocols as the RNA-Seq data. Isolation of high-purity smRNA/miRNA was performed using the Sigma-Aldrich mirPremier™ microRNA Isolation Kit (No. SNC50, Sigma-Aldrich, USA) following the manufacturer's suggested protocol for plant tissues. In order to maximize the capture of smRNA/miRNA, on-column DNase I digestion and ethanol precipitation is not recommended, thus, was not performed. Quantification of smRNA sample quality and concentration was performed using the Experion 'StdSens' kit (Bio-Rad Laboratories, Inc., Hercules, CA). Only one smRNA library (CAXXN), representing the male $F_2$ progeny individual, 11X-317-108, failed qPCR, and was not sequenced. Otherwise, 47 single-end (1×76) smRNA libraries were sequenced using an Illumina NextSeq system.

Small RNA reads were mapped to the full v5.1 reference genome containing the full chr15W and chr15Z assemblies using the program ShortStack to identify small RNA loci and putative miRNAs. miRNA loci are relatively small, can align to non-genic regions, and alignment quality can be affected by as little as one nucleotide, therefore necessitating use of the full chr15Z chromosome during alignment rather than the reduced chr15Z utilized in RNA-Seq mapping. The online database RNA Central was used to compare predicted miRNAs to homologous miRNAs in other species (rnacentral.org). psRNATarget (plantgrn.noble.org/psRNATarget) was used to identify putative target sites of predicted miRNAs in the *Salix purpurea* genome. DESeq2 was used calculate the sex-based differential expression of the predicted miRNAs using the same parameters as conducted with the RNA-Seq data.

*Bisulfite Sequencing*

For bisulfite sequencing, six male and six female $F_2$ progeny, for which mRNA-Seq and smRNA-Seq had also been generated, along with twelve males and twelve females from a diverse population of unrelated S. purpurea collected from across Europe and North America, were collected in April 2020 using the same protocols as the RNA-Seq data. Isolation of whole DNA was done using a Qiagen Plant mini kit following the recommended protocol. Paired end (1x151) libraries were generated using the Illumina NovaSeq 6000 system. Genomic DNA was first sheared to ~ 500bp using sonication (Covaris LE220), then subject to end repair, A-tailing and ligation of Methylated Indexed Illumina Adaptor (IDT). Bisulphite conversion and clean-up of adapter ligated DNA was done using Zymo EZ DNA Methylation-Lightening Kit. Final Bisulphite converted library was amplified using 10 cycles PCR and purified using AMPure Purification.

Raw fastq file reads were filtered and trimmed using the JGI QC pipeline resulting in the filtered fastq file. Using BBDuk (https://sourceforge.net/projects/bbmap/), raw reads were evaluated for artifact sequence by kmer matching (kmer=23), allowing 1 mismatch and detected artifact was trimmed from the 3' end of the reads. Spike-in reads, PhiX reads and reads containing any Ns were removed. Quality trimming was performed using the phred trimming method set at Q6. Following trimming, reads under the length threshold were removed (minimum length 25 bases or 1/3 of the original read length - whichever is longer). Finally, one base off the right of the reads were trimmed to prevent creation completely contained read pairs. Read mapping and methylation calling was done using Bismark. For read mapping, bowtie1 was used with seed length set to 50, maximum mismatches in the seed set to 1 and maximum insert size set to 1000. Methylation detection was performed using the bismark_methylation_extractor tool from the Bismark package using default settings except to ignore 10 bases of both the 5' and 3' ends of read1 and read2 to eliminate any bias seen in the ends of the reads. Tiling of methylated sites into regions and calling of differentially methylated regions was done using DMRfinder.

Gentotyping-by-Sequencing

SNP information was obtained using genotyping-by-sequencing (GBS) for each sample in the $F_2$ population, along with the parents and grandparents. GBS reads were mapped to the v5.1 reference genome using BWA-MEM, and TASSEL 5.0 was used to call variants. In order to call SNPs across the SDR, two mapping files were created per individual, each including the autosomes plus one of the sex chromosomes. This ensured independent and accurate SNP calling across the length of each sex chromosome. SNPs in the SDR, therefore, represent homologous loci between chr15Z, which can freely recombine with another 15Z homolog in males, and chr15W, which shows suppressed recombination across the SDR. The resulting VCF files were merged across loci with identical names into a single SNP file containing separate variant calls for 15Z and 15W. SNP information was filtered to remove loci that occurred in fewer than 90% of samples or with a minor allele frequency < 0.05. Following imputation and error correction, SNPs were phased to the grandparent haplotypes using TASSEL 5.0, resulting in 8,806 SNPs after filtering.
